# Statistical inference with a manifold-constrained RNA velocity model uncovers cell cycle speed modulations

**DOI:** 10.1101/2024.01.18.576093

**Authors:** Alex R. Lederer, Maxine Leonardi, Lorenzo Talamanca, Antonio Herrera, Colas Droin, Irina Khven, Hugo J.F. Carvalho, Alessandro Valente, Albert Dominguez Mantes, Pau Mulet Arabí, Luca Pinello, Felix Naef, Gioele La Manno

## Abstract

Across a range of biological processes, cells undergo coordinated changes in gene expression, resulting in transcriptome dynamics that unfold within a low-dimensional manifold. Single-cell RNA-sequencing (scRNA-seq) only measures temporal snapshots of gene expression. However, information on the underlying low-dimensional dynamics can be extracted using RNA velocity, which models unspliced and spliced RNA abundances to estimate the rate of change of gene expression. Available RNA velocity algorithms can be fragile and rely on heuristics that lack statistical control. Moreover, the estimated vector field is not dynamically consistent with the traversed gene expression manifold. Here, we develop a generative model of RNA velocity and a Bayesian inference approach that solves these problems. Our model couples velocity field and manifold estimation in a reformulated, unified framework, so as to coherently identify the parameters of an autonomous dynamical system. Focusing on the cell cycle, we implemented *VeloCycle* to study gene regulation dynamics on one-dimensional periodic manifolds and validated using live-imaging its ability to infer actual cell cycle periods. We benchmarked RNA velocity inference with sensitivity analyses and demonstrated one- and multiple-sample testing. We also conducted Markov chain Monte Carlo inference on the model, uncovering key relationships between gene-specific kinetics and our gene-independent velocity estimate. Finally, we applied *VeloCycle* to *in vivo* samples and *in vitro* genome-wide Perturb-seq, revealing regionally-defined proliferation modes in neural progenitors and the effect of gene knockdowns on cell cycle speed. Ultimately, *VeloCycle* expands the scRNA-seq analysis toolkit with a modular and statistically rigorous RNA velocity inference framework.

## INTRODUCTION

Single-cell RNA-sequencing (scRNA-seq) captures a static snapshot of gene expression in a destructive manner, making it difficult to interpret dynamical aspects of biological processes. To address this issue, computational approaches have emerged that reconstruct temporal information among cellular states from scRNA-seq data [1]. For example, RNA velocity exploits the ratio between unspliced and spliced transcripts to estimate a vector that describes the rate of change of gene expression [2]. The model considers a system of first-order ordinary differential equations describing the mRNA life cycle and whose key parameters are splicing and degradation rates. Under simplified assumptions, it is possible to estimate these parameters from data [3].

The original RNA velocity framework, implemented in *velocyto*, fixes a common splicing rate across genes to infer a relative gene-dependent degradation rate from spliced-unspliced phase portraits [2]. This parameter is then plugged into the differential equations to obtain a gene-specific velocity. An extended model for the estimation of RNA velocity is the “dynamical model,” implemented for the first time in the tool *scvelo*, which introduced for each gene a cell-wise latent time to support the estimation of kinetic parameters varying across a pseudotemporal axis, making them directly identifiable [4]. By exploiting expectation-maximization, *scvelo* estimates latent time and kinetic parameters. Other methods have harnessed these modeling ideas or worked towards extending them [5–14]. However, RNA velocity analysis remains highly sensitive to preprocessing choices and requires various heuristics to obtain the final estimates.

A pervasive yet potentially dangerous heuristic is the nearest-neighbor smoothing used to approximate expectations on the RNA counts; this procedure can let information bleed from some genes to others and cause distortions [15]. Additionally, the use of general non-linear dimensionality reduction techniques to bring the high dimensional velocity vector onto a two-dimensional embedding (e.g., UMAP, tSNE) risks introducing artifacts [16]. For instance, velocities associated with orthogonal processes, such as proliferation and differentiation, may be blended together, and adjacent yet unrelated cell populations might affect the resulting vector. Other algorithmic steps and corner cases that typically require attention have already been noted [2,17]. However, a seldom discussed, yet central, limitation of most RNA velocity models is that velocity estimation is not performed jointly on all genes. This strategy is problematic, even when some form of global reconciliation is sought; for example, when aggregating individual latent times into a global one, the obtained kinetic parameter estimates remain independent. This leads to a physically and geometrically inconsistent velocity vector, whose gene-specific components are on different timescales and whose resulting direction is not necessarily tangent to the low dimensional manifold cells traverse. This is inappropriate for unbiased forecasting, as future states predicted by integration are bound to rapidly escape the gene manifold and inhabit unlikely regions of the expression space.

Finally, the lack of established ground truths for RNA velocity limits the rigorousness of sensitivity analyses that can be performed on newly developed methods, creating a challenging environment to benchmark advanced extensions [18–21]. In particular, overparameterization becomes a concern, especially for models with less stringent assumptions, several non-linearities, or many degrees of freedom. Furthermore, proposed Bayesian formulations of the “dynamical model” return a high-dimensional mean-field posterior, which is not consistent with the assumption of low rank dynamics and is poorly suited to inference on the velocity and statistical comparisons of cell population dynamics.

We addressed these challenges by reformulating RNA velocity analysis as an inferential framework rooted in a manifold-constrained probabilistic model. Adopting this approach, we propose an explicit parametrization of RNA velocity as a field defined on the manifold coordinates. We focus on one-dimensional periodic manifolds in a framework called *VeloCycle*, enabling model validation and application to cell cycle dynamics. The cell cycle is the most ubiquitous periodic process in biology and plays a fundamental role in embryonic development, tissue regeneration, and disease [22,23]. Despite being pervasive in scRNA-seq datasets, default cell cycle analysis pipelines [24,25] are still restricted to categorical phase assignment based on a small selection of marker genes [26–28]. In this work, we not only tackle the broader issue of maintaining geometrical constraints during velocity estimation, but also make strides in improving cell cycle analysis in scRNA-seq data, highlighting its continuous nature and providing control over the actual biological time scales. We apply *VeloCycle* across different biological contexts, experimentally benchmark against time-lapse microscopy measurements, and illustrate the ability to perform statistical tests.

## RESULTS

### Manifold-constrained RNA velocity addresses shortcomings of other approaches

We first sought to redesign RNA velocity estimation by unifying manifold and velocity inference into a single probabilistic framework (**Fig. 1a, left**). This framework is articulated around a generative model with explicit low-dimensional dynamics at its core. In our model, cells move in time as points on a low-dimensional manifold *x* embedded within the space of all measured genes. Spliced and unspliced molecules are formulated as a function of *x* only (i.e. *s(x), u(x)*). Then, by parameterizing velocity as a function of the manifold coordinates *V(x)*, we constrain RNA velocity vectors to lie tangent to the manifold (**Fig. 1a, right**). This is contrary to previous approaches where velocity direction is unconstrained, as it is the result of gene-wise estimates [15,17] (**Fig. 1b**). We take the derivative of the expected spliced counts, apply the chain rule, and plug in the kinetic equations to obtain a velocity vector field interlocking the kinetic parameters of all genes and the dynamics of the latent coordinates (**see Methods 1-3**). Noise in the measured raw read counts is modeled as a negative binomial, also as a function of the manifold *x*, and biochemically informed priors are chosen for all other parameters, including splicing (β) and degradation (γ) rates for each gene (**Fig. 1c; see Methods 4**).

**Figure 1.**
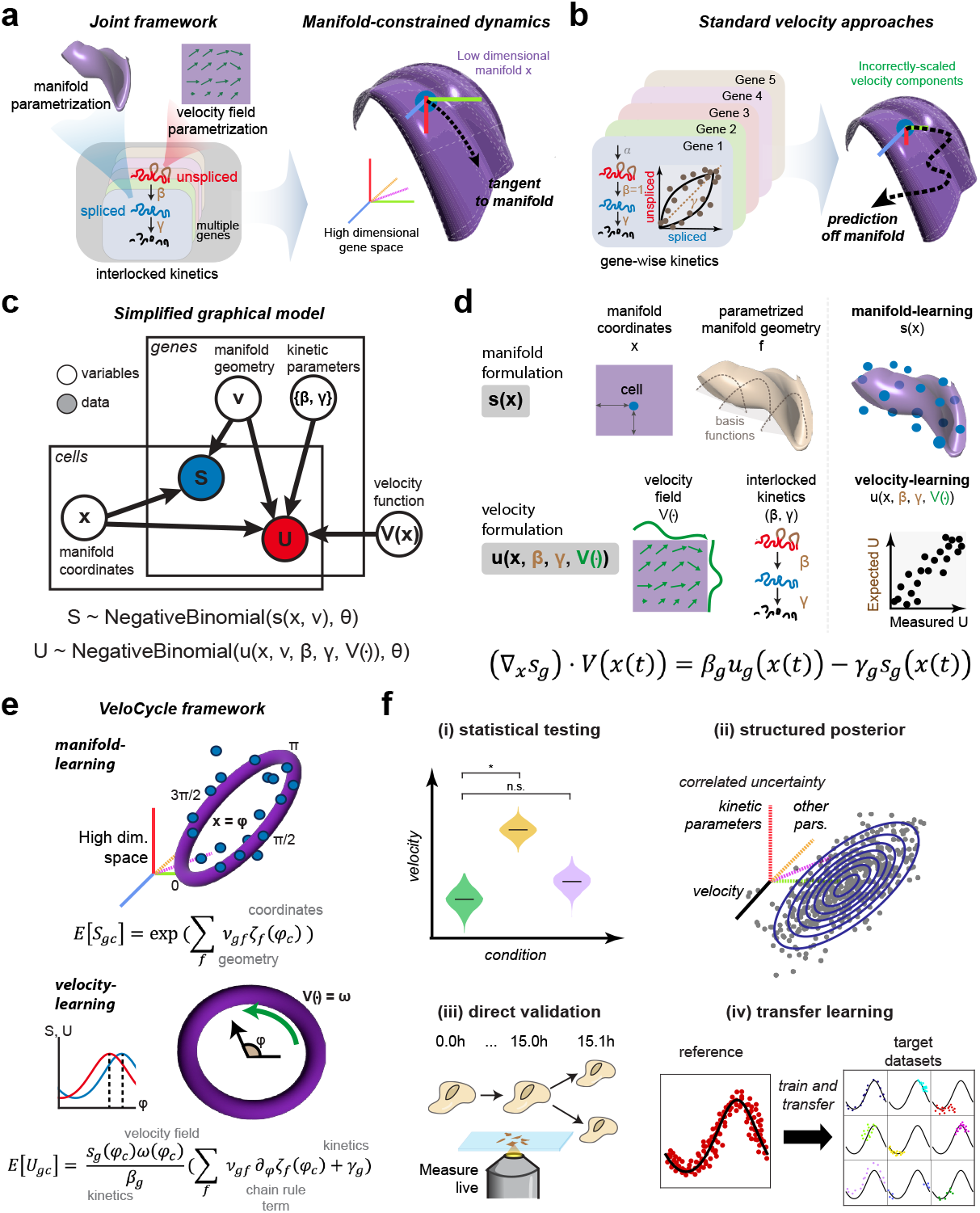
Statistical inference of RNA velocity with a manifold-constrained framework for the cell cycle. **(a)** Schematic of a joint framework for parameterization of the gene expression manifold and RNA velocity field. By defining velocity as a function of the manifold coordinates, the velocity vector field is constrained to be tangent to the manifold. This is achieved by interlocking the kinetic parameters of all genes with latent coordinate dynamics. **(b)** Schematic of unconstrained velocity estimation described by standard velocity approaches. By estimating the vector field as a combination of incorrectly-scaled, gene-dependent components, velocity is no longer tangent to the manifold. **(c)** Plate diagram of the probabilistic relationship among latent variables and observable data (S, U), modeled using a negative binomial distribution. S is sampled from the expectation, manifold coordinates, and manifold geometry. U is sampled from the manifold information, kinetic parameters, and velocity function. **(d)** Top: manifold formulation is defined for the spliced counts (s) using the cell-specific manifold coordinates (x) and a gene-specific geometric family (f), with which observed data can be directly mapped to the high-dimensional manifold space. Bottom: velocity formulation is defined for unspliced counts (u) as a velocity field function (V) and interlocked kinetic parameters (β, γ). We can obtain a velocity estimate by taking the chain rule over these entities. **(e)** Schematic of the two procedure steps used by *VeloCycle* to solve manifold-constrained velocity estimation for periodic processes such as the cell cycle. First, *manifold-learning* estimates the manifold coordinates and geometry; second, *velocity-learning* estimates the kinetic parameters and manifold-dependent velocity function. **(f)** Schematic of some types of velocity analyses that are possible for the first time with *VeloCycle*, including: (i) statistical credibility testing between multiple samples and against a zero-velocity null hypothesis; (ii) posterior marginal distribution analysis of velocity and kinetic parameters by Monte-Carlo Markov Chain (MCMC) sampling; (iii) extrapolation of velocity to real biological time of cell cycle speed with live microscopy; and (iv) transfer learning of gene manifold from high-content, quality references to low-content, noisy target datasets.

This formulation constitutes a latent variable framework for estimation of the gene expression manifold and RNA velocity. The choice of a specific dimensionality, topology, and associated functional parametrization constraining its geometry can be tailored in an application-specific manner (**Fig. 1d**). We propose inference in two statistical learning procedures: (1) *manifold-learning* to jointly learn the parameters defining the geometry of the gene expression space and assign each cell a manifold (latent) coordinate, and (2) *velocity-learning* to find a velocity field and kinetic parameters, conditioned on the manifold geometry and cell coordinates (**Fig. 1d-e**).

We implemented this scheme considering a scenario where the prior information on manifold topology is strong: the cell cycle, a one-dimensional periodic space on which gene expression varies smoothly and can be parametrized using a Fourier series. Our framework, *VeloCycle*, constitutes a generative probabilistic model with two groups of latent variables and is solved in Pyro [29] (**see Methods 4, Table 1**). The first group relates to manifold-learning and defines the low-dimensional manifold *x* parameterized as cell cycle phase (φ) and gene-specific Fourier coefficients (ν_0_, ν_1sin_, ν_1cos_) using the expected spliced counts as a function of the phase (**Fig. 1e and S1a-b**). The second group relates to velocity-learning from the expected unspliced counts and includes the gene-specific degradation rates (γ_g_), effective splicing rates (β_g_) and velocity harmonic coefficients (νω), which parameterize an angular speed function (ω(φ)) describing how cell cycle velocity changes along the manifold (φ) (**Fig. 1e and S1c-d; see Methods 4**). Using stochastic variational inference (SVI), *VeloCycle* returns the joint posterior probability of the latent variables, which can be used to (i) perform statistical velocity significance testing, (ii) characterize underlying correlations between the uncertainty of latent variables, (iii) estimate cell cycle velocities on a biologically-relevant time scale, and (iv) facilitate the application of velocity to small datasets by transfer learning (**Fig. 1f**).

**Table 1.** Overview of VeloCycle latent variables.

| Variable | Description | General Name | Training Step | Dimensions |
| --- | --- | --- | --- | --- |
| $\varphi_{xy}$ | cell cycle phase | Manifold coordinates | manifold-learning | (cell) |
| $v$ | Fourier coefficients for the genes | Manifold geometry | manifold-learning | (gene, harmonics) |
| $\Delta v$ | batch-specific expression offset | Data-specific noise in manifold geometry | manifold-learning | (batch, gene) |
| $shape_{inv}$ | spliced negative binomial noise | Measurement noise | manifold-learning | (gene) |
| $\log \beta_g$ | log splicing rate | Velocity kinetics | velocity-learning | (gene) |
| $\log \gamma_g$ | log degradation rate | Velocity kinetics | velocity-learning | (gene) |
| $v\omega$ | Fourier coefficients for the angular speed | Velocity function | velocity-learning | (condition, harmonics) |

### Sensitivity analysis on simulated data validates *VeloCycle*

After designing our model, we sought to evaluate its performance on simulated data, as no real dataset is endowed with ground truth information for phases, speed, and RNA kinetic parameters. We employed a simulation intended to preserve important relations expected in real data [2] and avoid biologically improbable scenarios (**see Methods 5; S2a-c**). Specifically, we incorporated positive correlations among the splicing and degradation rates (r=0.30) and baseline expression levels (r=0.30) (**Fig. S2a**). This structure naturally imposed a positive correlation between the splicing rate and total spliced counts as well as a negative correlation between the splicing rate and total unspliced counts (**Fig. S2b-c**).

First, we evaluated *manifold-learning* across 20 individually simulated datasets each containing 3,000 cells and 300 genes and found *VeloCycle* inferred phases that closely matched the ground truth, with a circular correlation of r_φ_ = 0.95 (**Fig. 2a-b**). The estimation error was consistently smaller than the uncertainty defined by the posterior, with true values falling within the 5%-95% credible interval for 99.2% of cells (**Fig. S2d**). We also verified that the gene-specific Fourier series coefficients closely tracked the original ground truths (r_ν0_ = 0.95, r_ν1sin_ = 0.98, and r_ν1cos_ = 0.98) (**Fig. 2c and S2e**). For these parameters, wider credible intervals corresponded to more noisy genes with a larger coefficient of variation (**Fig. S2f**). Overall, these results confirmed that *VeloCycle* correctly identified the manifold geometry and cell coordinates. To assess robustness of the model on different dataset sizes, we performed sensitivity analysis, varying the number of cells and genes (**see Methods 5**). We found that estimates were broadly accurate, with a circular correlation coefficient greater than 0.70 obtained using as few as 100 cells or 100 genes (**Fig. 2d**). We further benchmarked our inference against DeepCycle, a recent autoencoder-based method [30]. This comparison showed that *VeloCycle* was typically more accurate (60% lower MSE on average, rφ = 0.95) than DeepCycle (rφ = 0.73), despite the latter using velocity moments to achieve its estimations (**Fig. 2e-h**).

**Figure 2.**
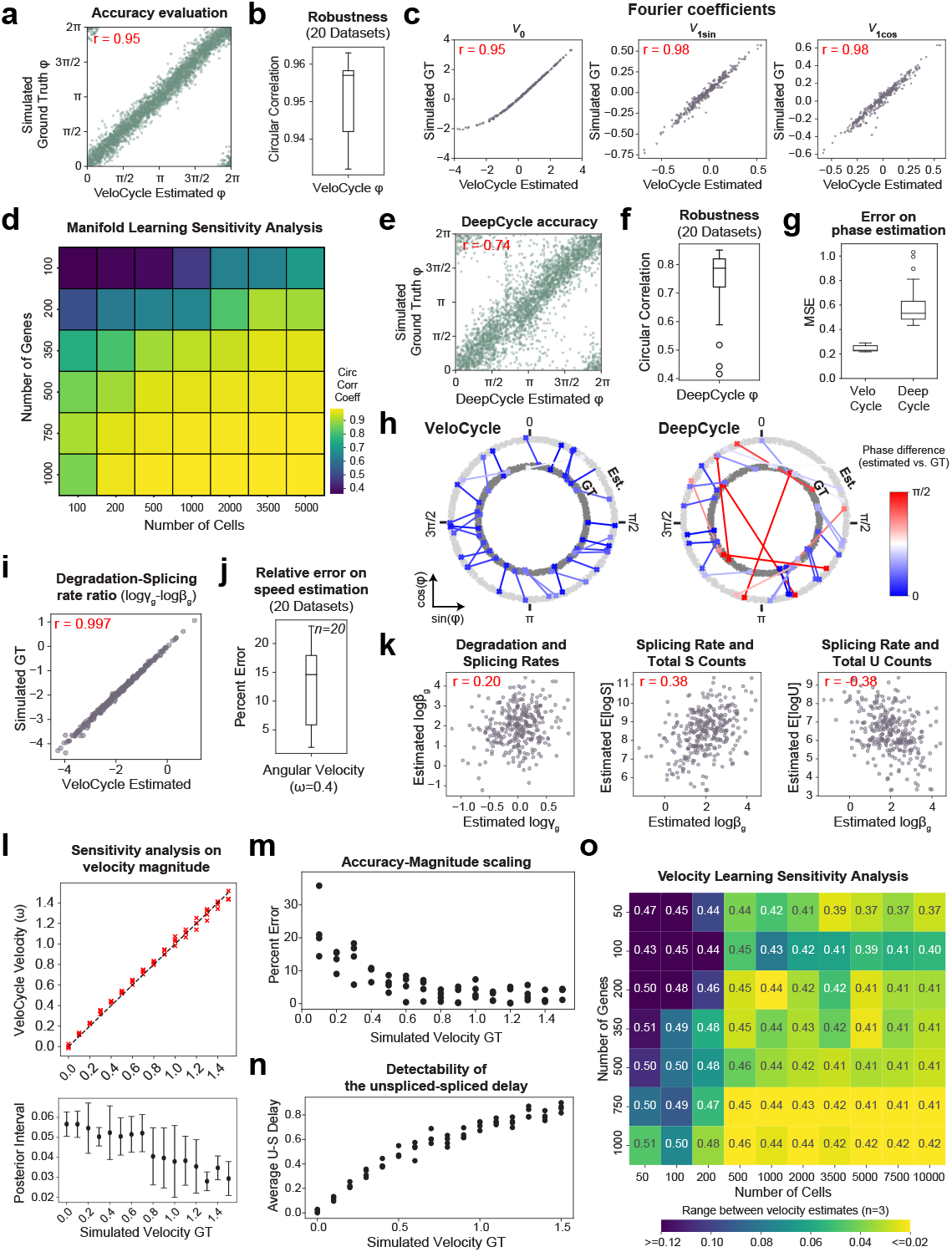
Sensitivity analysis of *VeloCycle* on simulated data. **(a)** Scatter plot of cell cycle phase assignment (VeloCycle Estimated) compared to the simulated ground truth (GT). **(b)** Box plot of circular correlation coefficients between estimated and GT phases across 20 independently simulated datasets, each containing 3,000 cells and 300 genes. **(c)** Scatter plots of estimated and GT values for the gene harmonic coefficients (v_0_, v_1sin_, v_1cos_) using the dataset in (a). **(d)** Heatmap of the mean circular correlation coefficient between estimated and GT phases computed with varying numbers of cells and genes. Each value is the average of three independent simulations. **(e)** Scatter plot of cell cycle phase estimation obtained by DeepCycle compared to simulated ground truth. The same dataset shown in (a) was used. **(f)** Box plot of circular correlation coefficients between DeepCycle-estimated and GT phases across the datasets shown in (b). **(g)** Box plots of per-cell mean squared error (MSE) for phase estimation with *VeloCycle* and DeepCycle. **(h)** Polar plots representing the phase difference between estimated and simulated GT for 30 randomly chosen cells from a single simulated dataset using *VeloCycle* (left) and DeepCycle (right). Each dot represents a cell, and lines connect the estimated phase assignment (Est; light gray) to simulated ground truth (GT; dark gray). **(i)** Scatter plot of estimated ratio between γ_g_ and β_g_ compared to simulated GT for 300 genes. **(j)** Box plot of percent error between estimated and GT velocity (ω) across 20 simulated datasets with a GT of 0.4. **(k)** Scatter plots illustrating the recovered relationships among splicing rate (logβ_g_), degradation rate (logγ_g_), spliced counts, and unspliced counts for 300 simulated genes. **(l)** Top: scatter plot of estimated and GT estimates for 16 different simulated velocities between 0.0 to 1.5 radians per mean half-life (rpmh) for 4 independently-simulated datasets. Bottom: box plots of posterior uncertainty intervals corresponding to the above simulations. **(m)** Scatter plot of percent error between estimated and GT velocity across conditions in (l). **(n)** Scatter plot of mean unspliced-spliced expression delay across conditions in (l). **(o)** Sensitivity analysis heatmap of the range among velocity estimates for 3 independently-simulated datasets, using varying numbers of cells and genes. The text value in each box represents the mean velocity over the 3 datasets, and intensity of the heatmap represents absolute range. The Pearson’s correlation coefficient (r) over 20 individual simulated datasets is indicated in red in (a), (c), (e), (i), and (k). Each green dot represents a single gene in (a) and (e). Each purple dot represents a single gene in (c), (i), and (k).

Next, we conditioned *VeloCycle* on the simulated phase and gene harmonics to assess *velocity-learning*. We observed accurate estimation of gene-wise kinetic parameters across 20 individually simulated datasets, with a particularly close match of degradation-splicing rate ratios to the ground truth (r_γ/β_ = 0.997, r_β_ = 0.918, r_γ_ = 0.617; **Fig. 2i and S2g-h**). Importantly, *VeloCycle* was capable of returning an accurate estimate of the mean angular velocity (percent error running 5.4-22.6%; **Fig. 2j**). *VeloCycle* recovered the biological correlation structure among estimated kinetic parameters and total counts, without imposing them in the model formulation (**Fig. 2k, cf. Fig. S2a-b**).

We performed sensitivity analysis to understand how the estimations behaved at different ground truth velocities. We considered a large span of cell cycle velocities fully encompassing the range of biologically plausible ones (16 values from 0 to 1.5 radians per mean half-life, or rpmh, four simulations each). The results highlighted a stable performance of the method, with estimates 0.2-35.8% away from the ground truth (**Fig. 2l-m**). Error increased at slower velocities, with a lower Pearson’s correlation between kinetic parameters and ground truths (**Fig. S2i, left**). Indeed, slower velocities corresponded to shorter delays between unspliced and spliced RNAs (**Fig. 2n; see Methods 4.5**), which are more difficult to characterize accurately. In all simulations, the degradation-splicing rate ratios almost perfectly matched the ground truth (mean r_γ/β_=0.99) (**Fig. S2i, right**). Finally, we investigated whether *velocity-learning* performance was affected by dataset size. We detected a dependence on the number of cells and genes, with the highest accuracy and tightest posterior ranges obtained on larger datasets; however, using more cells could compensate for fewer genes, and vice versa (**Fig. 2o and S2j**). We established 500 cells (and a minimum of 50 genes) or 350 genes (and a minimum of 50 cells) as the lower limits at which accurate velocity estimation can be performed.

### *VeloCycle* manifold-learning estimates accurate and robust phases

After validating on simulated data, we deployed *VeloCycle* on real datasets produced with different scRNA-seq chemistries. We reasoned that access to a cell cycle phase ground truth, even if categorical (e.g., G1, S, G2/M), would facilitate the evaluation of our phase assignments. Thus, we performed *manifold-learning* on a Smart-seq2 dataset of FUCCI system-transduced mouse embryonic stem cells (mESC) that were index-sorted using fluorescence-activated cell sorting (FACS) [31]. We fit the cell cycle phase on spliced counts using a gene set representing a broad gene ontology (GO) query [32] (**see Methods 4.3, 6.1**) and evaluated the results against FUCCI-FACS categories. Cells belonging to the same category were assigned to similar phases (**Fig. 3a-b**); a classifier based on two thresholds and trained on *VeloCycle* phases achieved 82.7% accuracy in predicting the annotations, almost matching the 87.8% accuracy obtained when training a logistic classifier on all genes (**Fig. 3c**). Furthermore, gene fits underlying *manifold-learning* closely replicated the expected sequential patterns of cell cycle genes. Among fits of high confidence were early-peaking histone acetylase *Hat1*, followed by transcription factor *Trp53*, and the later anaphase-promoting complex member *Ube2c* (**Fig. 3d**). The gene succession and oscillation amplitude were recapitulated when performing *manifold-learning* on a smaller set of 209 genes, anticipating that the method is effective on chemistries with lower sensitivity (**Fig. 3e-f**).

**Figure 3.**
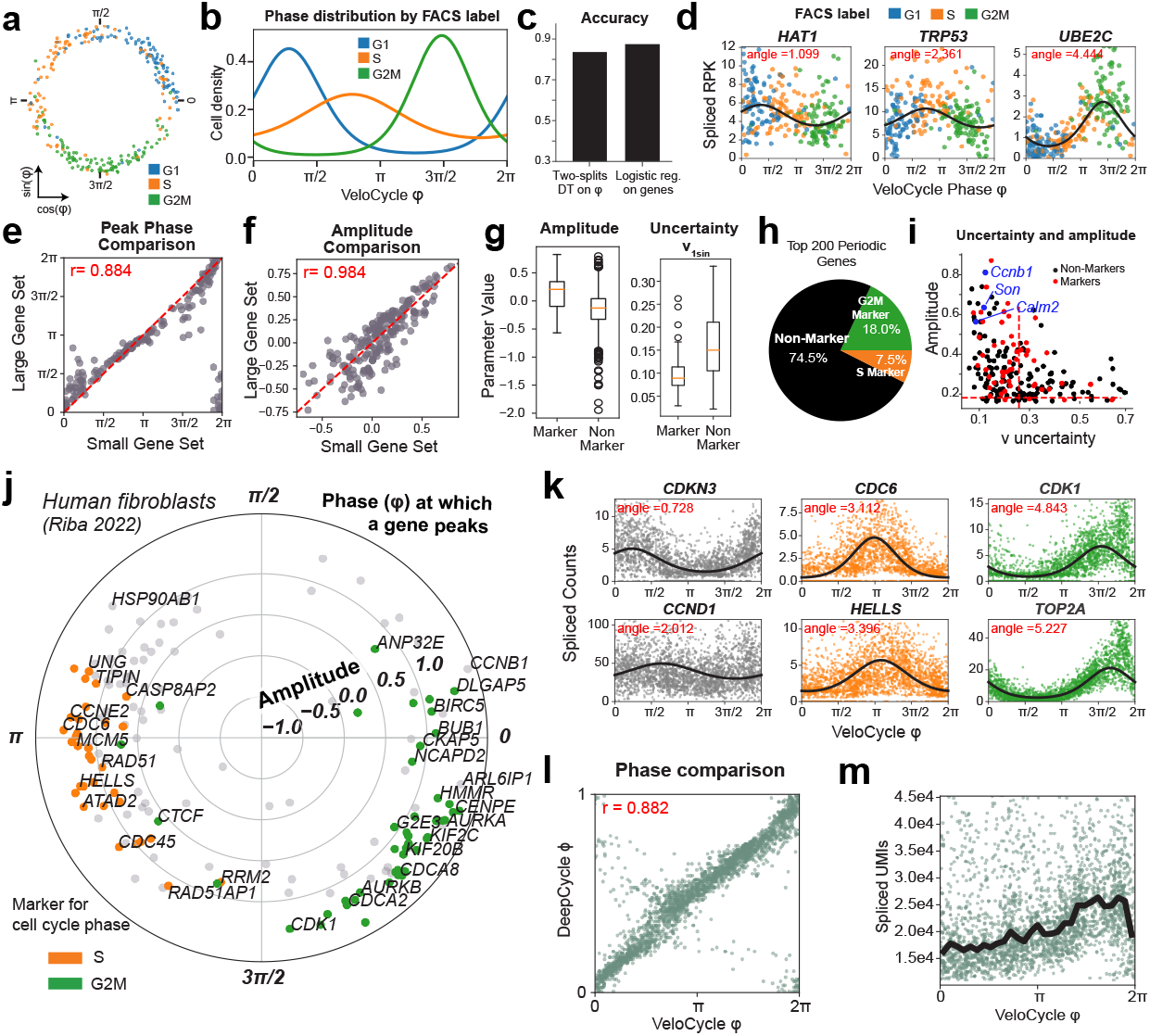
*Manifold-learning* and gene periodicity on different datasets and technolgies. **(a)** Scatter plot representing the phase assigment of 279 mouse embryonic stem cells, colored by their FACS-sorted categorical phase (G1, S, G2/M) [31]. **(b)** Density plot for FACS-sorted labels (G2, S, G2M) across the phase assigned by *VeloCycle*. **(c)** Bar plot reporting categorical phase predictor obtained using a two-thresholds decision tree trained on the *VeloCycle* phase estimates only versus a logistic regression classifier trained on the entire gene expression matrix. **(d)** Representative scatter plots of genes fits. Curved black lines indicate a gene-specific Fourier series obtained with *manifold-learning*. The “peak” indicates the position of maximum expression along the cell cycle manifold (VeloCycle φ). **(e)** Scatter plot of gene-wise peak position of maximum expression using a small (x-axis) or large (y-axis) gene set during *manifold-learning* for FACS-sorted mESC data. **(f)** Scatter plot of gene-wise peak position of gene-wise amplitude using a small or large gene set. **(g)** Box plots of gene-wise amplitude and harmonic coefficient uncertainties for marker and non-marker genes for FACS-sorted mESC. **(h)** Pie chart of categorical composition for the top 200 periodic genes, as determined by amplitude. **(i)** Scatter plot of gene-wise total harmonic coefficient (ν) uncertainty and amplitude. Gene dots are colored as standard “markers” or “non-markers”. Red dashed lines represent the mean values for “markers.” **(j)** Polar plot of estimated gene harmonics for human fibroblasts data [30]. Each dot represents a gene (n=160). The position along the circle represents the phase of maximum expression, and the distance from the center represents total amplitude. Colored genes (orange/green) are those used to compute a standard cell cycle score with *scanpy* or Seurat [24,25]. **(k)** Selected scatter plots of genes fits for markers of early (*CDKN3, CCND1*), mid (*CDC6, HELLS*), and late (*CDK1, TOP2A*) cell cycle progression obtained for the human fibroblasts data. **(l)** Scatter plot of phases estimated with *VeloCycle* compared to DeepCycle for 2,557 human fibroblasts. The circular correlation is indicated in red. **(m)** Scatter plot of total raw spliced UMI counts by *VeloCycle* phase. Black lines indicate the binned mean UMI level.

Given our Fourier parametrization, we could classify genes by the phase of peak expression, oscillation amplitude, and estimation uncertainty (**Table S1**). Inspection of phase-amplitude relations revealed that marker genes typically used for scoring in packages such as Seurat and scanpy [24,25] (henceforth “standard markers”) clustered by phase, consistent with the FACS-based ground truth (**Fig. 3g-h**). Compared to non-markers, standard markers on average had a higher amplitude (mean 0.14 versus -0.15) and lower posterior uncertainty (standard deviation 0.26 versus 0.43) (**Fig. 3g**). However, of the top 200 periodic genes based on amplitude, the vast majority (74.5%) were not standard markers (**Fig. 3h**), and many (n=78) could be equally or more confidently trusted (i.e., tighter posterior probability) as cell phase predictors (**Fig. 3i**). Among those were calcium-binding protein *Calm2*, splicing co-factor *Son*, and cyclin *Ccnb1*, which all play roles in cell proliferation [33–36].

We continued our scrutiny of *manifold-learning* using 10X Chromium data of human fibroblasts (**Fig. 3j-k and Table S2**). To put *VeloCycle* in relation to other approaches, we compared its estimated phases to those obtained by DeepCycle [30], finding a strong correspondence (human fibroblasts: r=0.882; **Fig. 3l**). Therefore, *VeloCycle* accomplishes similar phase estimation to DeepCycle but without using velocity and in tandem with fitting individual gene harmonics. As further validation that the correct cell cycle dynamics were captured, we observed a gradual increase in total UMIs along the phase, followed by a sharp drop corresponding to cytoplasm partitioning during cytokinesis (**Fig. 3m**). These results highlight that *manifold-learning* estimates a biologically-meaningful one-dimensional geometric space that tracks with the cell cycle across scRNA-seq chemistries.

### Unspliced-spliced delays along *VeloCycle* phase identify realistic cell cycle velocities

We next investigated whether the unspliced molecule counts together with the *VeloCycle* phase are sufficiently informative to estimate cell-cycle velocity. To explore this intuitively before performing the full inference, one can extract phases and gene harmonics with *manifold-learning* for unspliced and spliced UMIs independently and use an approximate formula for the velocity that we derived (**see Methods 4.2, 6.7, 7**). We applied this approach on two cultures of human RPE1 cells that were grown in parallel and under identical conditions so that we could also assess robustness by replicate comparison. First, we extracted the phases on each of the datasets by *manifold-learning*, then we measured the delays (i.e. the phase difference) between peak unspliced and spliced expression for each gene (**Fig. 4a**). We observed consistent and positive delays for the genes (**Fig. 4b**) that correlated well between replicates (r=0.90; **Fig. 4c**). We interpreted this correlation as the first evidence that the data contains velocity information on the cell cycle, so we proceeded to estimate a cell cycle period with the aforementioned approximate formula. The calculation returned a period 18.5 times the average half-life, which corresponds to 18.5h assuming a realistic average half-life of 1h (**Fig. 4d**). In addition to being an approximation, another limitation of the point estimate is that it is not based on a proper noise model and is not associated with an uncertainty measure. To obtain a more accurate estimate and statistical measures of confidence, we learned the complete Bayesian model (*velocity-learning*) on both RPE1 replicates, conditioning on the random variables inferred by *manifold-learning*. Scaling the obtained velocity by the fitted average half-lives yielded average cell cycle periods of 20.1h ± 0.2h and 20.0h ± 0.2h (mean ± 95% credible intervals) for the two replicates (**Fig. 4e**). The posterior distributions broadly overlapped (71.2% overlap), indicating no credible velocity difference between the two replicates. To confirm on real data that *VeloCycle* can estimate cell cycle speed along a dynamic range relevant biologically, we performed *velocity-learning* on mESC, a rapidly-cycling cell type [37,38]. For this dataset, *VeloCycle* returned an estimation of 10.5 ± 0.3 average half-life (**Fig. S3a**). As with RPE1 cells, the model recovered kinetic parameters with expected relationships among total UMI counts and gene-specific splicing and degradation rates, as previously observed in simulated data (**Fig. S3b-e, cf. Fig. 2i**). Taken together, these findings confirm *VeloCycle* can estimate a cell cycle velocity and sample informative posterior distributions.

**Figure 4.**
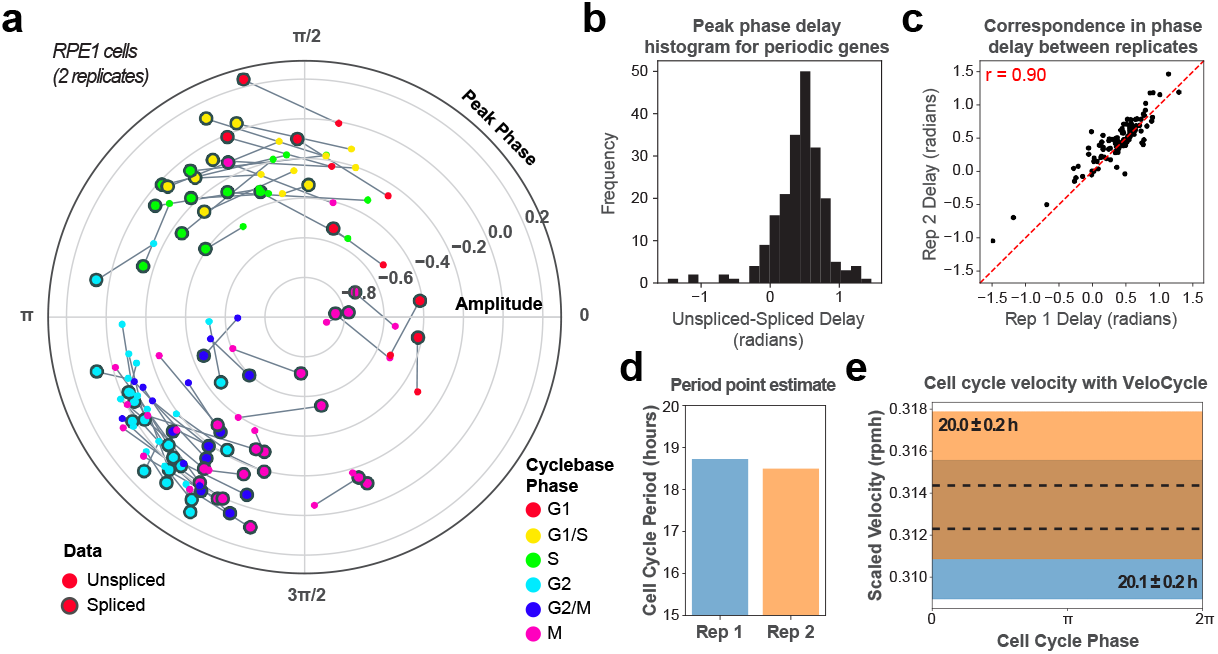
Analysis of delays and velocity scale in RPE1 cells. (**a**) Polar plot of the peak unspliced and spliced expression for 106 marker genes across two scRNA-seq replicates of RPE1 cells (4,265 and 9,994 cells) analyzed with *manifold-learning*. Genes are colored by their categorical annotation in Cyclebase 3.0 [67]. Unspliced gene fits were inferred separately, conditioned on cell phases obtained when running *manifold-learning* on the spliced UMIs. (**b**) Histogram of unspliced-spliced delays (in radians) for 106 genes. Pearson’s correlation is indicated in red. (**c**) Scatter plot of unspliced-spliced delays between two RPE1 cell line replicates. (**d**) Bar plot of the cell cycle periods obtained with a first-order-approximate point estimate (**see Methods**). (**e**) Posterior estimate plot of constant, scaled cell cycle speed (radians per mean half-life, rpmh) in two RPE1 cell line replicates. The black dashed lines indicate a mean of 500 posterior predictions, and the colored bar indicates the credibility interval (5th-95th percentile).

### A structured variational distribution preserves uncertainty correlations and leads to better uncertainty estimates

Although we showed our variational formulation recovers accurate estimates of cell cycle phase and velocity in simulated and real data using stochastic variational inference (SVI), it is reasonable to question the limits of a simplified mean-field variational family in representing the structure of joint uncertainty among latent variables. We hypothesized that such a parametrization choice may lead to an overconfidence in the estimated velocity posterior because uncertainties on these latent variables may be inherently correlated (**Fig. 5a**). A piece of evidence in this direction was the observation that estimates on random gene subsets fell outside the posterior credible interval of the fit on all genes (**Fig. 5b**). To eliminate this bias towards the underestimation of velocity uncertainty, we decided to characterize the model joint posterior by sampling it with Markov Chain Monte Carlo (MCMC; **see Methods 4**). Using a No U-Turn Sampler, we studied the posterior for human fibroblasts [30], with MCMC revealing a five-times wider uncertainty compared to mean-field SVI (0.10 rpmh vs 0.02 rpmh; **Fig. 5c**).

**Figure 5.**
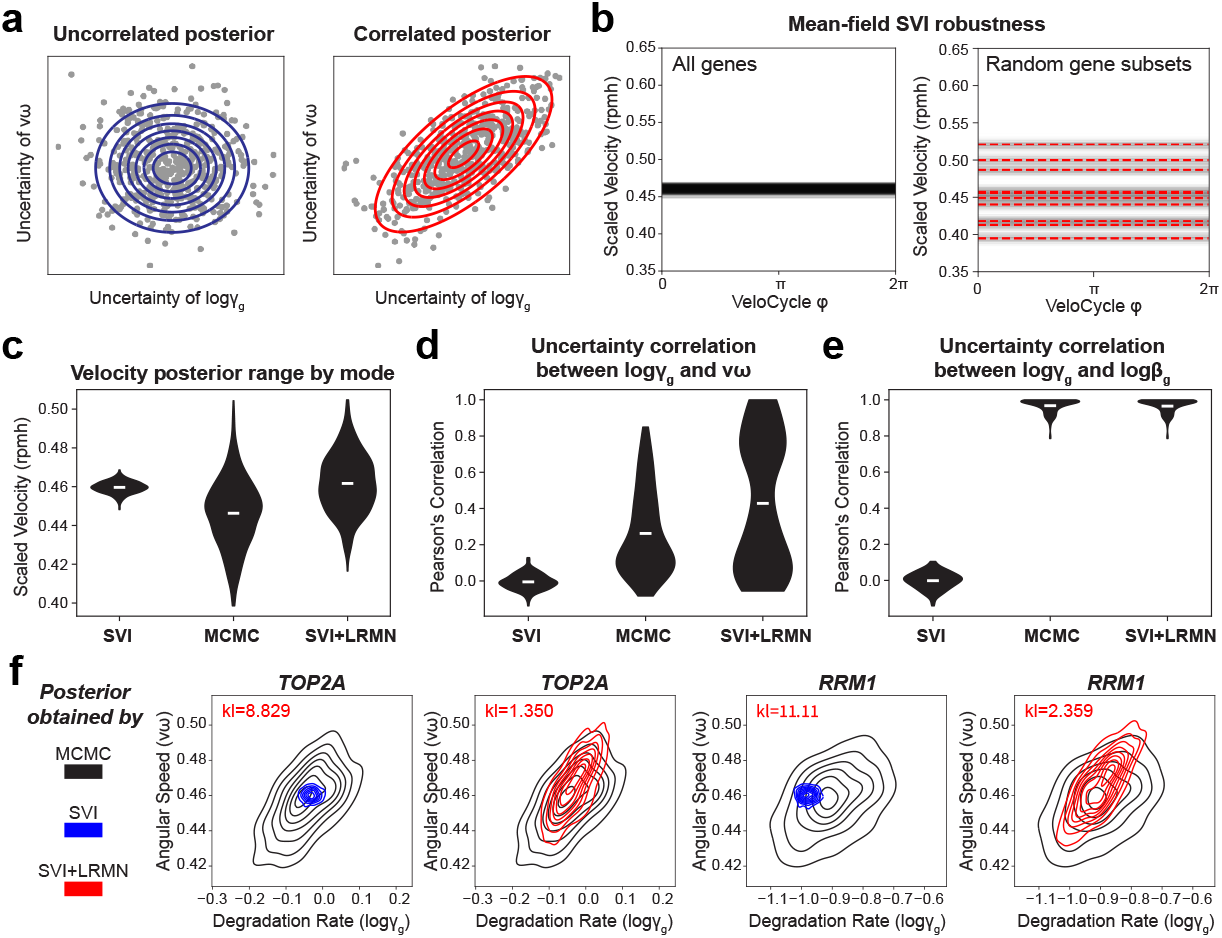
Relationships between parameter uncertainties and choice of the variational distribution. (**a**) Schematic of the hypothetical scenarios where a gene has uncorrelated (left) and correlated (right) posterior uncertainty between logγ_g_ and νω. Blue circles represent the Gaussian kernel density of the distribution, and red lines represent an uncertainty interval between two arbitrary fixed points. (**b**) Left: posterior estimated velocity plot inferred for 2,557 cultured human fibroblasts [30] using the original stochastic variational inference (SVI) mode of *VeloCycle*. Right: posterior estimated velocity plot on human fibroblasts, estimated with *VeloCycle* for ten random subsamples of the data, each using only 50% of the genes. (**c**) Violin plots of scaled velocity (in rpmh) for human fibroblasts after estimation using the stochastic variational inference (SVI), Monte Carlo Markov Chain (MCMC), and low rank multivariate normal (SVI+LRMN) *velocity-learning* modes. (**d**) Violin plots of Pearson’s correlations between the degradation rate (logγ_g_) and angular speed (νω) posterior uncertainties across 160 genes using different *VeloCycle* modes. (**e**) Violin plots of Pearson’s correlations between the degradation (logγ_g_) and splicing (logβ_g_) posterior uncertainties across 160 genes using different *VeloCycle* models. (**f**) Density representation of overlapping logγ_g_-νω posterior distributions between MCMC and either SVI (top) or SVI+LRMN (bottom) for *TOP2A* and *RRM2* (black: MCMC; blue: SVI; red: SVI+LRMN). Kullback-Leibler divergence scores are shown in red. All posterior means were taken over 500 predictive samples.

Consistent with our hypothesis, this wider credible interval manifested along with a correlated joint posterior, capturing dependencies among the uncertainty of different latent variables. Specifically, examining the posterior, we found samples of the angular speed (νω) and degradation rate (logγ_g_) for certain genes that exposed a correlation structure (mean r = 0.26; **Fig. 5d**). Moreover, for each gene we noticed a strong correlation (mean r = 0.96) between posterior samples of splicing (logβ_g_) and degradation (logγ_g_) rates (**Fig. 5e**). Both features cannot be captured by a mean-field variational distribution.

These findings advocated for a recrafting of our variational distribution to accommodate typical features of the posterior inferred by MCMC, in order to maintain inferential accuracy but avoid significantly time-consuming sampling procedures. We reformulated our variational distribution with logγ_g_ and νω modeled as a low rank multivariate normal (LRMN) and with the logβ for each gene modeled as a normal conditional on the corresponding logγ (**see Methods 3.2**). Upon retraining this new SVI+LRMN model, we obtained a velocity estimate with a larger uncertainty range (0.08 rpmh) than with mean-field SVI (**Fig. 5c-e**). Additionally, we detected a correlation among the SVI+LRMN posterior samples between logγg and νω for a subset of genes that overlapped with the results of MCMC; this resulted in a decreased Kullback-Leibler (KL) divergence between the SVI+LRMN and MCMC posteriors than between the SVI and MCMC posteriors (**Fig. 5f and S4a**).

Importantly, there was a correspondence between the specific genes with high logγ_g_ and νω uncertainty correlation in both SVI+LRMN and MCMC (**Fig. S4b**). Genes with a greater correlation between logγ_g_ and νω tended to be those with larger unspliced-spliced delay (**Fig. S4c**). We speculated the degree of dependence between a gene’s logγ_g_ and νω is related to the extent it contributes to the velocity estimate. This was supported by a leave-one-out experiment, where individual genes with smaller degradation rates were those most strongly affecting velocity estimates (**Fig. S4c-d**). The correlation between logγ_g_ and νω posterior uncertainty was also reproducible when SVI+LRMN was applied to mouse ESCs (**Fig. S4e-f**). Overall, these implementation changes led to generation of a more robust model that can be confidently used for inference, while preserving the underlying correlation structure of the true posterior.

### Cell tracking and labeling experiments validate computationally inferred velocities

Estimates of a manifold-constrained cell cycle speed with *VeloCycle* are most conveniently expressed in units of mean half-lives (**see Methods 4.1, 4.6**). Since the average values of half-lives are typically known in many cell types, real time estimates of RNA velocity can be obtained and validated along the cycle. In this respect, we reasoned that time-lapse microscopy offers a compelling means for comparing *VeloCycle* estimates to a ground truth.

To benchmark our velocity estimation framework against an experimentally-determined cell cycle period, we examined a dataset of dermal human fibroblasts (dHFs) monitored by time-lapse microscopy and for which scRNA-seq data was collected (**see Methods 6.3**) [39]. Our SVI+LRMN model inferred a constant cell cycle period of 15.3 ± 1.2h, assuming an average half-life of the modeled transcripts of 1h (**Fig. 6a and S5a-d**). Next, we used *VeloCycle* to infer a non-constant (periodic) cell cycle velocity, and we obtained a similar estimated duration of 16.5 ± 2.1h, with maximal velocity near mitosis (approximately 3π/2 < φ < 2π) (**Fig. 6b**). We then reconstructed the cell cycle period using cellpose and TrackMate for 268 individual cells followed by time-lapse imaging (**Fig. 6c**) [40,41]. From these data, we recovered a median cell cycle of 15.8h (s.d. 3.1h), which overlapped with the posterior credibility interval of the *VeloCycle* estimate (**Fig. 6d, c.f. Fig. 6a-b**). Comparable results were obtained when using the smaller set of cycling genes [30] (**Fig. S5e**). Taken together, these results indicate an ability to obtain comparable cell cycle speed estimates from live-microscopy and *VeloCycle*.

**Figure 6.**
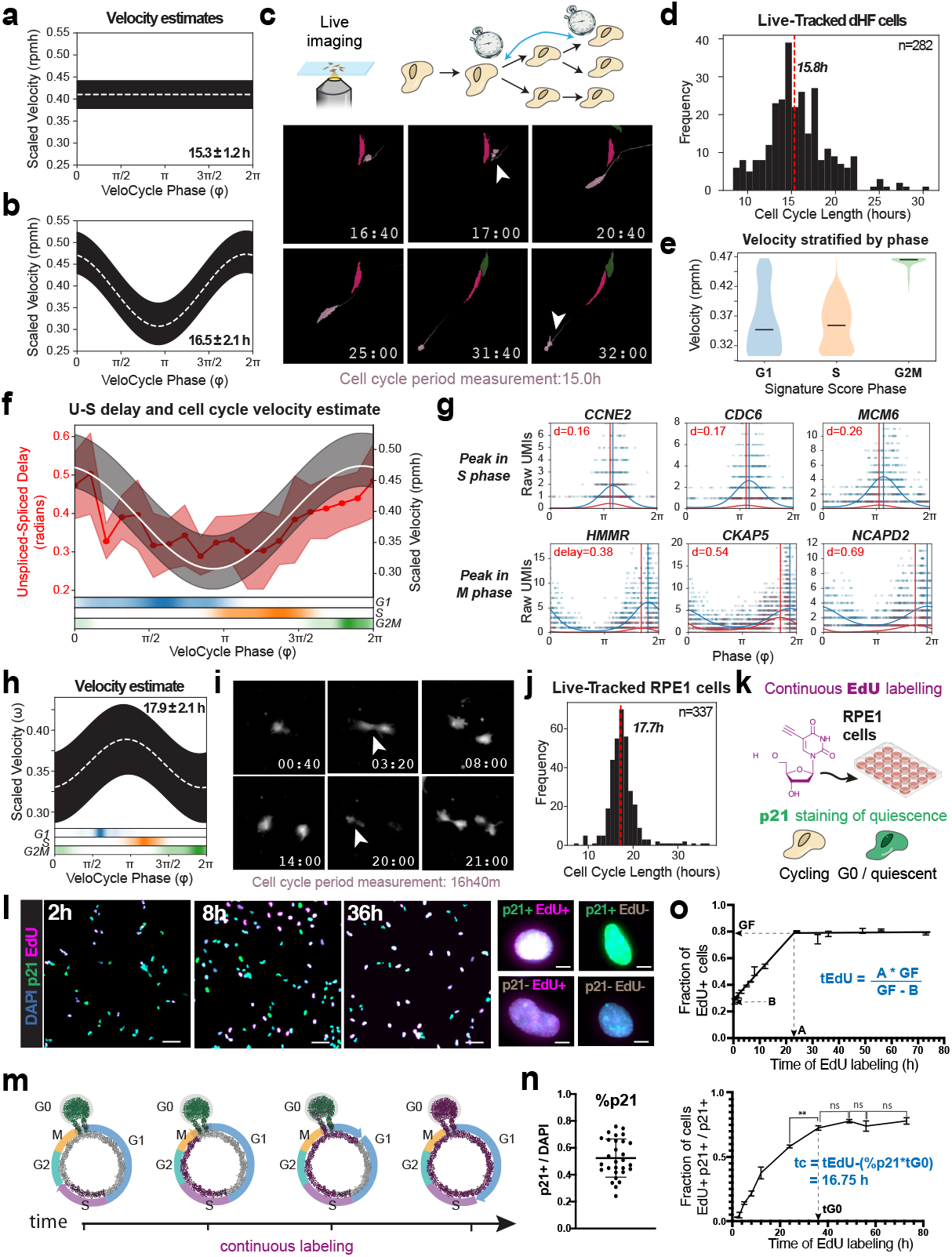
Validation of computationally inferred velocities by cell tracking and labeling experiments. **(a)** Posterior estimate plot of constant cell cycle speed in 1,222 dermal human fibroblasts (dHFs) [39]. **(b)** Posterior estimate plot of periodic (non-constant) cell cycle speed in cells from (a). **(c)** Top: schematic of time-lapse microscopy with live-imaging to track consecutive cell divisions. Bottom: example microscopy images at multiple time points to illustrate tracking a single segmented dHF cell (pink) through two divisions. Following division of the mother cell (16:40h), one daughter cell is tracked for 15 hours until dividing itself (31:40h). **(d)** Histogram of cell cycle period for 268 dHFs tracked by live-microscopy. **(e)** Violin plot of cell cycle speed for dHFs, stratified by categorical phase assignment. **(f)** Dual-axis plot of the correspondence between the unspliced-spliced expression delay (left) and cell cycle velocity estimate (right). Left: genes were grouped in 20 equal bins by phase and the unspliced-spliced delay was calculated. Right: the scaled velocity estimate from (c). Bottom: cell cycle categorical (G1, S, G2/M) phase assignment probability. **(g)** Gene expression scatter plots for genes peaking in S (top) and M (bottom) phases. Vertical lines correspond to the peak phase of spliced (blue) and unspliced (red) counts, used to compute an unspliced-spliced delay in (f). **(h)** Posterior estimate plot of periodic (non-constant) cell cycle speed in 3,354 retinal pigmented epithelial cells (RPE1). **(i)** Representative microscopy images tracking a single RPE1 cell from birth (3:20h) to subsequent division (20:00h). **(j)** Histogram of cell cycle period for 337 RPE1 cells tracked by live-microscopy. **(k)** Diagram of the cumulative EdU/p21 experiment. Cells were continuously exposed to EdU, fixed at different timepoints, and subjected to EdU detection and p21 immunostaining. **(l)** Left: representative images of p21 (green), DAPI (cyan), and EdU (magenta) staining after cumulative EdU labeling for 2h, 8h, and 36h. Scale bar is 100um. Right: representative images of individual cells with different staining combinations. Scale bar is 10um. **(m)** Schematic of cumulative EdU labeling during cell cycle progression. Cycling cells incorporate EdU (magenta) when they undergo DNA replication (S phase). Thus, the duration of the EdU pulse is directly proportional to the fraction of EdU-positive cells. G0/quiescent cells are represented in green. The total number of cells consists of the number of cells in all cell cycle phases and G0, so the total duration of the cell cycle must be accounted for while excluding quiescent cells. **(n)** Dot plot representing the average percentage of p21+ cells along the different time points. Error bar indicates the standard deviation (SD), and each dot represents the percentage of p21+ cells for a single replicate (n=29). **(o)** Top: line plot of the fraction of EdU+ cells after at 13 time points (from 30 min to 73h). Data show the mean of three replicates (except for 2h, which is from two), and error bars indicate the SD. Accumulation of EdU can be divided into a linear growth phase and a plateau phase. Using quantities derived from a linear fit of the growth phase, we can derive a formula for calculating the time for total EdU (tEdU) labeling. Bottom: line plot of fraction of EdU-positive cells among quiescent cells (p21+) plotted as a function of time. The time when the fraction of EdU-positive cells among the quiescent population stops growing significantly is taken as an estimation of the tG0. The red dashed line indicates the median in (d) and (j). The white dashed line indicates the mean of 500 posterior predictions and the black bar indicates the credibility interval (5th-95th percentile) in (a), (b), and (h).

We next stratified velocity by an independent categorical cell cycle phase to gain further granularity on these evaluations and model behavior. We observed a faster progression through the cell cycle during G2/M phase (mean scaled velocity of 0.47 rpmh) compared to a slower progression during G1 (0.37 rpmh) and S (0.36 rpmh) phases (**Fig. 6e**). Kinetic parameters and their posterior uncertainties were strongly correlated between the constant and periodic velocity models (**Fig. S5f-g**). Interestingly, when estimating the average unspliced-spliced delay for genes peaking at different cell cycle phases, we found that cell cycle phases with larger average delays corresponded to regions with faster velocity (**Fig. 6f**). Genes with larger delays were also those with smaller splicing and degradation rates, which is expected from the approximate model (**Fig. S5h; see Methods 4.2**). After examining the unspliced-spliced delay and the low-rank gene-wise posterior correlation between the angular speed and degradation rate, we could identify specific genes that most strongly contributed to the underlying velocity estimates (**Fig. 6g**).

To further scrutinize the degree to which cell cycle durations inferred by *VeloCycle* match those obtained experimentally, we performed time-lapse microscopy and scRNA-seq on the same cultured RPE1 cells. The speed obtained with *VeloCycle* was approximately 17.7 ± 2.1h (**Fig. 6h, see Methods 7.3**); as in dHFs, this computational estimate overlapped with the mean cell cycle duration of 17.7h (standard deviation of 3.4h) obtained from tracking dividing cells by time-lapse imaging (338 cells) (**Fig. 6i-j**). We next sought to compare our cell cycle duration measurements from time-lapse microscopy and *VeloCycle* to those obtained using an orthogonal experimental technique. Therefore, we performed continuous EdU labeling to independently estimate cell cycle length (**Fig. 6k-l**). After monitoring EdU levels at 13 time points over 72 hours (**Fig. 6m**), we used p21 (CDKN1A) staining to account for cells in G0 and determined a mean cell cycle length of 16.8h (**Fig 6n-o; see Methods 7.4**). Taken together, these findings validated the computational RNA velocity estimates in the context of the cell cycle. To our knowledge, this is the first example of a direct validation of RNA velocity estimation with experimental methodologies and justifies the use of *VeloCycle* output in units of real (i.e., no pseudo-) time.

### *VeloCycle* enables direct statistical velocity comparisons in response to drug treatment

Existing frameworks for RNA velocity do not propose an approach to test the statistical significance of obtained estimates, likely because it is challenging given a gene-wise velocity parametrization. For example, it is currently not possible to determine whether RNA velocity estimates close to zero should just be interpreted as noise. Furthermore, direct comparisons between velocity estimates of two samples cannot be supported by a measure of confidence. With *VeloCycle*, statistical inference testing on velocity is possible for the first time, both against a specific null-hypothesis and for differential velocity significance between cell populations.

To illustrate how our model can be used for statistical velocity tests in practice, we conducted RNA velocity analysis on a PC9 adenocarcinoma cancer cell line before (D0) and after (D3) treatment with the drug erlotinib [42] (**Fig. S6a-e**). Statistical testing in a Bayesian setting can be achieved by calculating credible intervals from the posterior. First, we considered the velocity posterior of the untreated (D0) cells to ask whether there is statistical support for a non-zero velocity. Given no overlap between the credible interval and zero, we could conclude the data contains statistically significant evidence for progression through the cell cycle (**Fig. 7a, left**). We then compared the treated sample (D3), with the control (D0). We found significant velocity differences between the D0 and D3 time points, where a slower mitotic cell cycle speed was detected at D3 (**Fig. 7b**). Such testing can be done globally and also locally. For example, we stratified by phase intervals and inspected the posterior samples, confirming a decreased speed during G2/M phase at D3 compared to D0, but not during G1 and S (**Fig. 7b and Fig. S6f**). The reduced presence of cells in M phase after erlotinib treatment was further suggested by the low density of D3 cells with assigned the phase coordinate (**Fig. S6a, bottom**).

**Figure 7.**
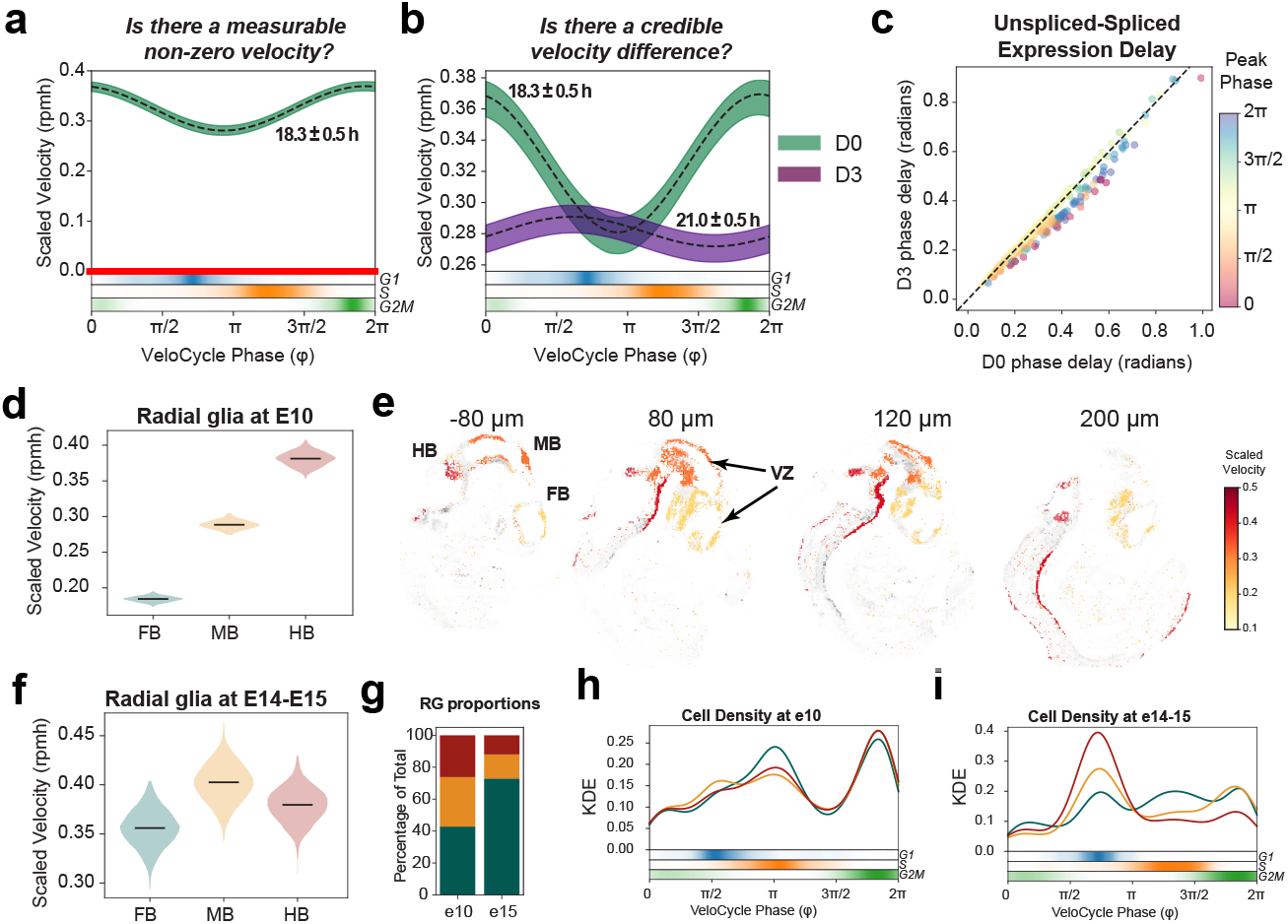
*VeloCycle* statistical inference on lung adenocarcinoma and neural progenitors. **(a)** Posterior estimate plot of scaled velocity in the PC9 lung adenocarcinoma cell line (D0: 9,927 cells) compared to a zero-velocity control (in red) [42]. **(b)** Posterior estimate plot of scaled velocity before (D0) and after (D3: 3,943 cells) PC9 treatment with erlotinib. Dashed lines indicate the mean of 500 posterior predictions; green (D0) and purple (D3) bars represent credibility intervals (5th-95th percentile). Areas in which the intervals do not overlap indicate statistically significant velocity differences. Bottom: cell cycle categorical (G1, S, G2/M) phase assignment probabilities. **(c)** Scatter plot of the mean unspliced-spliced expression delay for 273 genes between D0 and D3 samples. Gene dots are colored by peak expression phase. **(d)** Violin plots of scaled velocity estimates obtained for mouse forebrain (FB: 3,293 cells), midbrain (MB: 2,388 cells), and hindbrain (HB: 2,012 cells) radial glial (RG) progenitors at developmental stage E10 [48]. Black dashed lines indicate the mean of 500 posterior predictions. **(e)** Spatial projection of single cell clusters onto four sections of a reference E11 mouse embryo profiled with spatial transcriptomics (hybridization-based in situ sequencing; HybISS), colored by scaled velocity estimates obtained with *VeloCycle*. Regional domains (FB, MB, HB) and the ventricular zone (VZ) are labeled accordingly. Mapping of single cells to spatial data was achieved using the BoneFight algorithm [48]. **(f)** Violin plots of scaled velocity estimates for similar populations as on left (FB: 2,460 cells; MD: 307 cells; HB: 176 cells) at developmental stages E14/E15. **(g)** Bar plot of regional proportions of radial glia progenitor cells analyzed at early (E10) and late (E14/E15) time points. **(h)** Kernel density estimation plots of cell distributions along the cell cycle manifold at E10, colored by regional identity. **(i)** Kernel density estimation plots of cell distributions along the cell cycle manifold at E14/E15.

Since the unspliced-spliced delay is linked with cell cycle velocity, we hypothesized there would be differential delays between the D0 and D3 time points, particularly for genes peaking during M phase. After calculating the gene-wise unspliced-spliced delay before and after erlotinib treatment, we indeed noticed a subset of genes with peak expression during M phase and larger phase delays in D0 than D3 (**Fig. 7c**); this included anaphase-promoting complex member *CDC27* (differential delay, dd=0.11 radians), cyclin-dependent kinase inhibitor *CDKN3* (dd=0.10), and centrosome scaffolding factor *ODF2* (dd=0.09) (**Fig. S6g**). A decreased cell cycle speed specifically during M phase is consistent with the expected effect of erlotinib, an EGF-blocker inhibiting progression to G1 [43]. The result also aligns with evidence that a complete arrest should not be observed for the PC9 cell line, which has been reported to have some resistance to a complete blockade [44–46].

### Cell cycle speed in radial glial progenitors varies along a spatio-temporal axis in mouse development

Regulation of proliferation rate as well as of symmetric and asymmetric divisions of radial glia cells (RG) in the ventricular-zone plays a critical role in controlled developmental timing along an anterior-posterior axis of the brain [47]. To elucidate whether there are differences in cell cycle speed among progenitors populating different spatial regions during mouse neurodevelopment, we performed *VeloCycle* estimation on forebrain (FB), midbrain (MB), and hindbrain (HB) RG at the embryonic day 10 (E10) stage [48]. Cell cycle speed varied along the forebrain-midbrain-hindbrain axis, with progenitors dividing more quickly posteriorly (HB) than anteriorly (FB) (**Fig. 7d**). A finer visualization of this gradient was allowed by computationally mapping the cell cycle speed inferred in these cells to the corresponding locations using in situ hybridization spatial transcriptomics (HybISS) data and the BoneFight algorithm [48] (**see Methods 6.5**). We observed rapidly dividing RG localized close to the ventricular zones, highlighting that cell proliferation takes place along the ventricular zone and suggesting that different segments of the zone proliferate at different rates (**Fig. 7e**) [49]. Conversely, at E14 and E15 time points, RG from all three brain regions stabilized at a similar proliferation speed, with no credible velocity difference (**Fig. 7f**). At these later time points, the majority of RG in the midbrain and hindbrain regions had accumulated in a non-proliferative state; the majority of RG cells present were from the forebrain, which more slowly developed at E10 (**Fig. 7g-i**). These results align with recent studies showing that hindbrain specifies into non-proliferating, differentiated cell types more quickly; an increased proliferative capacity is thus likely required in the earlier stages of development [50–52]. Furthermore the later slowdown is expected and in line with what has been reported in EdU tracking studies [53,54].

### Transfer learning of manifold parameters enables discovery of velocity alterations in genome-wide perturbation screens

Previous frameworks for RNA velocity have offered restricted applicability to samples containing few cells. With recent single cell technologies designed to screen the effects of hundreds of small genetic, environmental, or drug perturbations, there is a growing need to assess changes in cell dynamics under circumstances with limited data [55–57].

*VeloCycle*, with its manifold-constrained velocity estimates, can explore RNA velocity in such contexts: by transferring *manifold-learning* from a large dataset onto smaller datasets, one can perform velocity inference using limited cells or using cells representing only a portion of the phase space (**see Methods 6.6**). To demonstrate this, we studied a large-scale, genome-wide Perturb-seq dataset where hundreds of individual gene knockouts were introduced into the RPE1 cell line via a targeted, pooled CRISPR library, followed by scRNA-seq after seven days in culture [58]. First, we ran *VeloCycle* on non-targeted control cells (NT) and a pooled group of gene knockout conditions corresponding to well-characterized marker genes for the cell cycle (CC-KO). The cell cycle period was 25.6 ± 1.3 hours for NT and 30.9 ± 1.3 hours for CC-KO (**Fig. 8a**). Similar results were obtained when using a large set of cell cycle genes (n=426) compared to a smaller gene set (n=120) (**Fig. S7a-e**). When CC-KO conditions were stratified by genes typically considered S and G2/M markers, we observed an accumulation of cells in the G1 phase space compared to NT cells (**Fig. 8b and S7f**). This suggests the loss of function for some individual cell cycle related genes disrupts cell cycle progression, either by slowing down the proliferation rate in certain phases, or by halting progression altogether ahead of specific entry checkpoints.

**Figure 8.**
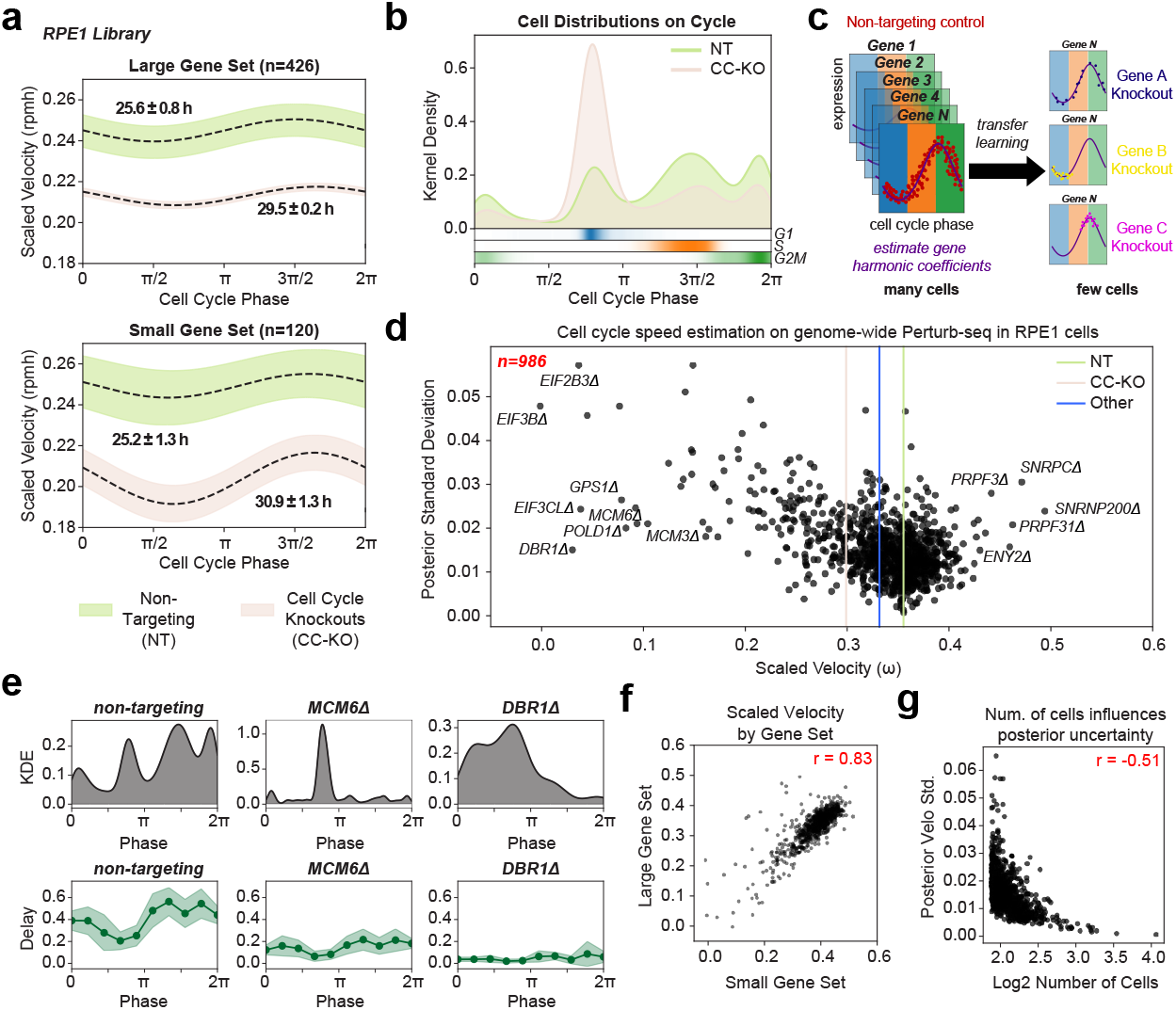
Transfer learning of manifold parameters to study effects of genome-wide knockouts on cell cycle velocity. **(a)** Posterior estimate plot of cell cycle speed for RPE1 cells 7 days after CRISPR-induced single-gene knockdowns with Perturb-seq, stratified by non-targeting controls (green; 11,485 cells) and cell cycle knockout (beige; 6,275 cells) conditions [58]. Manifold-learning was performed using either a large (top; n=426) or small (bottom; n=120) gene set. Black dashed lines represent the mean estimate of 500 posterior predictions. **(b)** Kernel density plot of continuous cell cycle phase distributions for non-targeting (NT) and cell cycle knockout (CC-KO) samples from (a). Heatmap bar plot (bottom) is of cell density by categorical phase assignment. **(c)** Schematic of the employed transfer learning approach. Gene harmonic coefficients are obtained on NT controls (with many cells) using the manifold-learning model, and are then applied to assign phases to cells with one of hundreds of gene knockout conditions (each with a few cells) potentially distributed unequally on the manifold. **(d)** Scatter plot of cell cycle *velocity-learning* estimates and posterior standard divisions for 986 individual cell cycle knockout (Δ) conditions in 167,119 RPE1 cells. Vertical lines correspond to the mean velocity estimates for non-targeting (green), cell cycle marker gene knockouts (tan) and other gene knockouts (blue). **(e)** Top: kernel density estimate (KDE) plots of cell distribution for the non-targeting, *MCM6Δ*, and *DBR1Δ* conditions. Bottom: binned unspliced-spliced expression delay (Delay) for the same conditions. Genes were binned by position of peak expression into 10 groups along the cell cycle to obtain an average delay. The dark green line represents the mean delay; the light green line represents the standard deviation. **(f)** Scatter plot of scaled cell cycle velocity estimates obtained for 986 conditions in (d) using small and large gene sets. **(g)** Scatter plot of total number of cells per condition and posterior velocity standard deviation for 986 conditions in (d). Pearson’s correlation coefficient is indicated in red in (f) and (g).

To scrutinize the effect of individual gene knockout conditions on cell cycle speed, we employed a transfer learning approach in which we conditioned our manifold-learning on gene harmonics previously inferred from the NT and CC-KO data subsets, assigning phases to a significantly larger population of 167,119 cells and 986 individual knockout conditions, some with as few as 75 cells (**Fig. 8c**). Consistent with coarser stratifications of the data, we observed a significant decrease in cell cycle speed in individual cell cycle related-gene knockout conditions compared to both non-targeting control cells and cells with gene knockdown unaffiliated with the cell cycle (**Fig. 8d**). Several of the most impaired cell cycle speeds were found in knockouts of highly characterized genes involved in DNA replication (*MCM3Δ* and *MCM6Δ*) and translation initiation (*E1F3BΔ, EIF2B3Δ, EIF3CLΔ*) (**Fig. 8d-e**). Curiously, knockout conditions for several splicing and mRNA processing genes either significantly decreased or increased the estimated cell cycle speed, including *DBR1Δ*, an intron-lariat splicing factor (11.7-fold decrease compared to NT condition), *PRPF3Δ* (1.2-fold increase), and *PRPF31Δ* (1.3-fold increase) (**Fig. 8d-e**). Given the dependence of RNA velocity estimation on the governing differential equations of the RNA metabolic life cycle, this result indicated that biological disruptions affiliated to RNA metabolism undermine the biophysical parameterization of the velocity framework. Moreover, the number of cells present in the dataset per condition had a direct influence on the velocity estimate posterior uncertainty, suggesting that more cells, and thereby less aggregated sparsity for a condition, increased the confidence of the *VeloCycle* model in the obtained velocity estimate **(Fig. 8f-g**). Ultimately, these analyses demonstrate that velocity can be applied, with transfer learning approaches, in large-scale perturbation contexts as a metric to assess the impact of gene knockouts on the dynamics of a biological process.

## Discussion

In this work, we address several limitations of current RNA velocity methods by designing a framework that unifies manifold and velocity inference into a single probabilistic generative model. We propose an explicit parametrization of RNA velocity as a vector field defined on the manifold coordinates, implemented and thoroughly tested for one dimensional periodic manifolds and the cell cycle (**see Methods 1-4**). RNA velocity has been previously applied to illustrate cell cycle progression, yet in ways that required several heuristics and with exclusive exploratory value, as no conclusion could be made from the inferential procedures [17,27,59,60].

*VeloCycle* uses variational inference to learn a Bayesian model that operates on raw data with appropriate noise models, instead of heuristic nearest-neighbor smoothing (**Fig. 1, see Methods 4**). *VeloCycle* returns uncertainty estimates, enabling direct evaluation of the confidence about the estimation results and cell cycle speed comparisons between samples. These capabilities are relevant in different biological settings, such as in cancer biology, where alterations to the cell cycle progression need to be scrutinized using snapshot single-cell data (**Fig. 7**). Therefore, *VeloCycle* could yield new biological insight into disease progression, for example by characterizing differences in proliferation rates between tumors across microenvironments or patients.

Uncertainty measurements are central to statistical evaluation of RNA velocity. The first methods to introduce Bayesian variational inference for RNA velocity modeling, VeloVAE [18], VeloVI [19], and Pyro-Velocity [20], simplify the variational distribution in ways that limit usefulness of the estimated joint posterior, particularly given an unscaled gene-wise velocity parametrization. More generally, models with a high number of degrees of freedom and the assumption of independence risk overfitting noise and overestimating confidence in the velocity [17,21]. In this study, we control for this risk by constraining all spliced-unspliced fits under a single velocity function, and we structure our model to scrutinize dependencies between the cell cycle velocity and kinetic parameters from the full posterior distribution (by MCMC and a LRMN variational distribution) (**Fig. 5, see Methods 3.2**). Moreover, we exploit the fact that this parameterization is explicit in the velocity to perform direct inference on a single latent variable.

While our model for RNA velocity estimation offers clear benefits, there remain open avenues for further development. First, while our mathematical framework is amenable to multidimensional formulations and various topologies, the current work focuses on the case of one-dimensional periodic manifolds. Thus, extensions of *VeloCycle* into higher-dimensional latent spaces can be naturally pursued, although significant efforts will be required to find appropriate parametrization of more complex manifolds. Second, the issue of defining dimensionality intersects with that of gene selection; different subspaces defined by unique genes expose distinct manifolds traversed by varying fractions of cells [17]. Methods developed with this problem in mind have been recently proposed [61], and with appropriate modifications, these could be integrated into RNA velocity estimation methods to automate topology and gene set selection. In this direction, frameworks that consider multiple manifolds with varying topologies, spanned by cells in different subspaces, while also assigning specific cells and genes to these features, will notably enhance the general applicability and utility of manifold-consistent RNA velocity estimation. Third, our model assumes a constant gene-specific splicing and degradation rate; in fact, for some genes, such rates likely change in different phases of the cell cycle [27,62]. A future extension to *VeloCycle* for which the kinetic parameters are defined by a parameterizable function, could address this limitation. Yet, maintaining the model well-conditioned in these settings might be non-trivial.

Widely used standard analysis pipelines use a small group of marker genes to attribute a categorical phase assignment to single cells, even though cell cycle progression is a continuous process [24,25]. Recent methods to infer continuous phase assignment represent a significant improvement over scoring-based approaches [63–66]. The *manifold-learning* of *VeloCycle* makes progress along this direction, also inferring individual gene periodicity patterns, providing posterior uncertainty and obtaining results that compare favorably with other methods [30] (**Fig. 3**). Importantly, the *manifold-learning* step is flexible and facilitates transfer learning: the geometry of the manifold can be estimated on a larger or higher quality dataset and serve as a prior for a smaller dataset. This enhances the robustness and applicability of velocity-learning across diverse experimental conditions. We employ these transfer learning capabilities on a Perturb-seq dataset, demonstrating that RNA velocity can be used as a readout of the context of a high-content screen (**Fig. 8**). This is particularly relevant given the increased use of barcoding strategies for single cell-level screening. We expect future applications of such models in the context of drug screening and evaluation of genetic changes on heterogeneous pools of cells.

A way to validate the overall consistency of a RNA velocity vector field has been to correlate an heursiticly estimated transition probability between populations with prior knowledge on their lineage relashionships; however, this is correlative and indirect [14]. Here, we instead compare directly estimates with the real velocity of the process. By specifically biologically-reasoned priors, velocities obtained with *VeloCycle* can be directly interpreted as the proliferation speed, which can vary in different tissue locations, at different moments of development, or as a result of perturbations to the core gene regulatory network [49,58]. This study empirically validated obtained RNA velocities, juxtaposing *VeloCycle* speed estimates with proliferation times obtained by live-cell microscopy imaging and cumulative EdU labeling (**Fig. 6**).

Ultimately, our framework represents an advancement in the rigor of dynamical estimations from single-cell data. The promising outcomes of tailoring RNA velocity to single processes advocates for the development of new models that dissect the high-dimensionality of single-cell data into individual biological axes with corresponding and interpretable RNA velocity fields.

## METHODS

Please see the attached document for the complete methods section.

## SUPPLEMENTAL TABLEs

**Table S1**. *VeloCycle* gene harmonic parameters for Smart-seq2 mESC obtained with *manifold-learning*

**Table S2**. *VeloCycle* gene harmonic parameters for 10X human fibroblasts obtained with *manifold-learning*

## DATA AVAILABILITY

The raw and processed scRNA-seq data in the RPE1 cell line that was newly generated for this study are available at GEO accession number **GSE250148**. All other scRNA-seq data used in this study were collected from previously published works [30,31,39,42,48,58] and relied on the cell type annotations made by the original authors. Jupyter notebooks and other affiliated files to reproduce the results shown in this study are provided via our GitHub page: https://github.com/lamanno-epfl/velocycle/. Processed versions of all published data (including spliced-unspliced counts matrices) are also available at the above link. The simulated scRNA-seq datasets, processed scRNA-seq metadata for the new RPE1 samples, cell tracking data from live-image microscopy and cumulative EdU staining experiments are also available at the above link..

## CODE AVAILABILITY

*VeloCycle* is implemented in Python and available as an open-source package on GitHub at https://github.com/lamanno-lab/velocycle. *VeloCycle* can be installed from PyPi using the command pip install velocycle or via direct installation from the GitHub page using the command *pip install git+https://github.com/lamanno-epfl/velocycle*. git@main. Python version 3.8 or newer is required. Source code, installation instructions, tutorials, and a file containing all required package dependencies are also available on GitHub. Additional code and notebooks to reproduce the results of this study are available via the link provided on GitHub.

## AUTHOR CONTRIBUTIONS

A.R.L. developed the idea, designed, implemented, and refined the model framework, analyzed the scRNA-seq and time-lapse microscopy data, created the figures, and wrote the manuscript. M.L., L.T., and C.D. participated in development of the idea and model formulation. A.H., A.V., I.K., and H.C. performed scRNA-seq experiments. A.H. and A.V. also conducted time-lapse microscopy and cumulative EdU experiments, with image processing analysis aid from A.D.M. P.M.A. helped test phase estimation approaches. L.P. helped refine the idea and related analyses. F.N. and G.L.M. developed the idea, supervised the project, and wrote the manuscript. All co-authors read and approved the manuscript.

## Acknowledgements

This project has been made possible in part thanks to Chan Zuckerberg Initiative grants number 2022-249212 and 2019-002427. G.M.L. received support from the Swiss National Science Foundation (SNSF) grant PZ00P3_193445. F.N. received support from the SNSF grant #310030B_201267 and the EPFL. L.P. is partially supported by the National Human Genome Research Institute (NHGRI) Genomic Innovator Award (R35HG010717). We thank members of the La Manno, Naef, Pinello, Williams and Castelo-Branco labs for their generous feedback and discussions on the project, particularly Daniil Bobrovskiy, Cameron Smith, Qian Qin, Eli Bingham, Luise Seeker, Nadine Bestard, Mukund Kabbe, and Fabio Baldivia Pohl. We also thank the entire EPFL Gene Expression Core Facility (GECF) for their assistance with scRNA-seq experiments.

## SUPPLEMENTAL FIGURES

**Figure S1.**
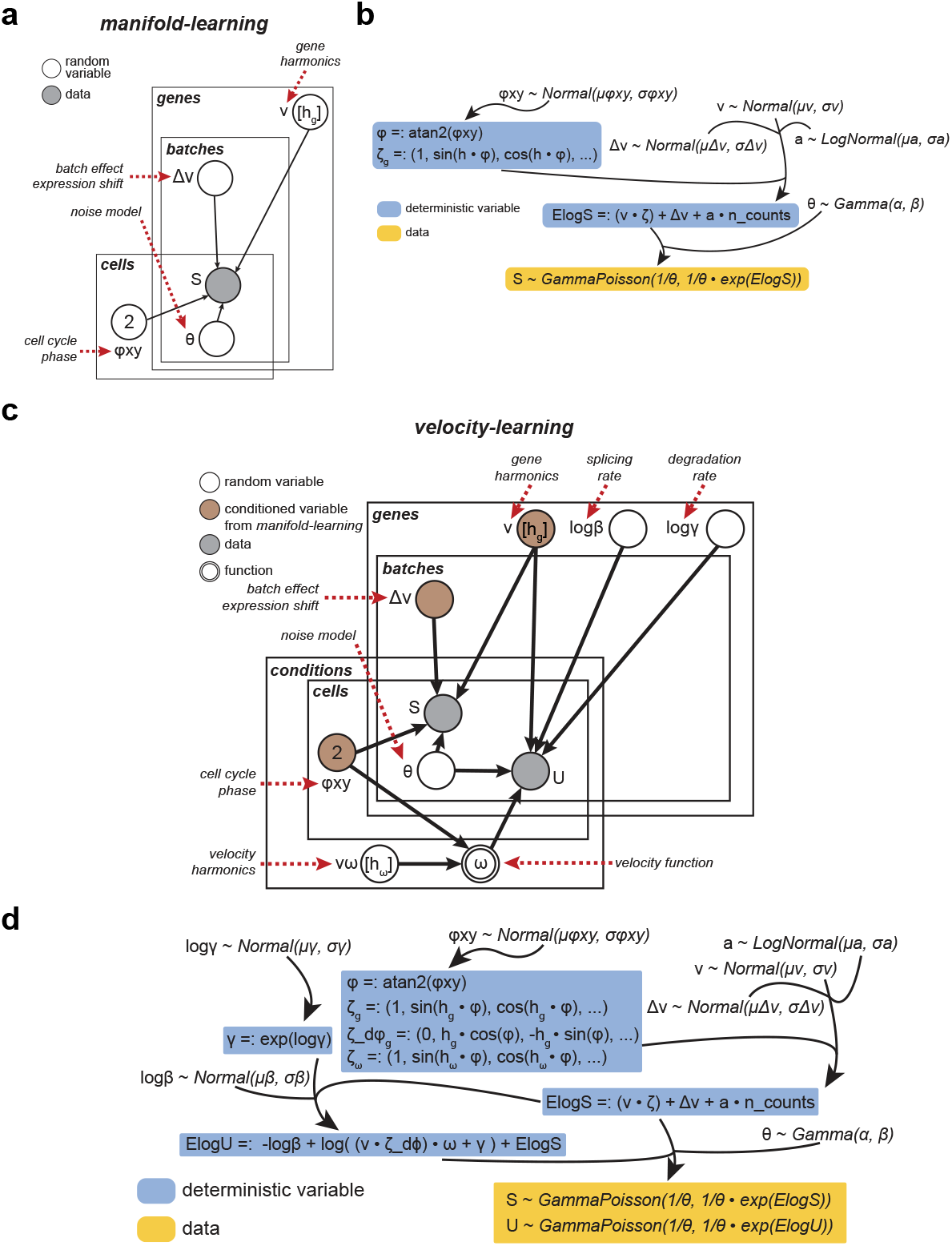
Plate diagram and mathematical formulation of *VeloCycle* framework for *manifold-learning* and *velocity-learning*. **(a)** Plate notation diagram of the *manifold-learning* procedure used to infer manifold coordinates (φ) and geometry (ν) given raw spliced count data. No k-nearest neighbor smoothing is performed. The model assigns each cell to a phase along the cell cycle (φ) and fits a set of harmonics (ν) for each individual gene using a Fourier series. **(b)** Formulaic representation of the *manifold-learning* procedure shown in (a). Spliced counts (S) are defined as the expectation (ElogS) plus noise, modeled after a negative binomial distribution. **(c)** Plate notation diagram of the complete *velocity-learning* procedure. Nodes indicate a model variable (white: random variable; gray: observed data; brown: conditioned variable from *manifold-learning*) and arrows indicate dependency. Each plate (genes, cells, conditions, and batches) signal independence and contain variables that are of the same dimensions. **(d)** Mathematical representation of the *velocity-learning* procedure shown in (c). Blue-boxed variables are deterministic and computed from latent variables; yellow-boxed variables are conditioned on observed data.

**Figure S2.**
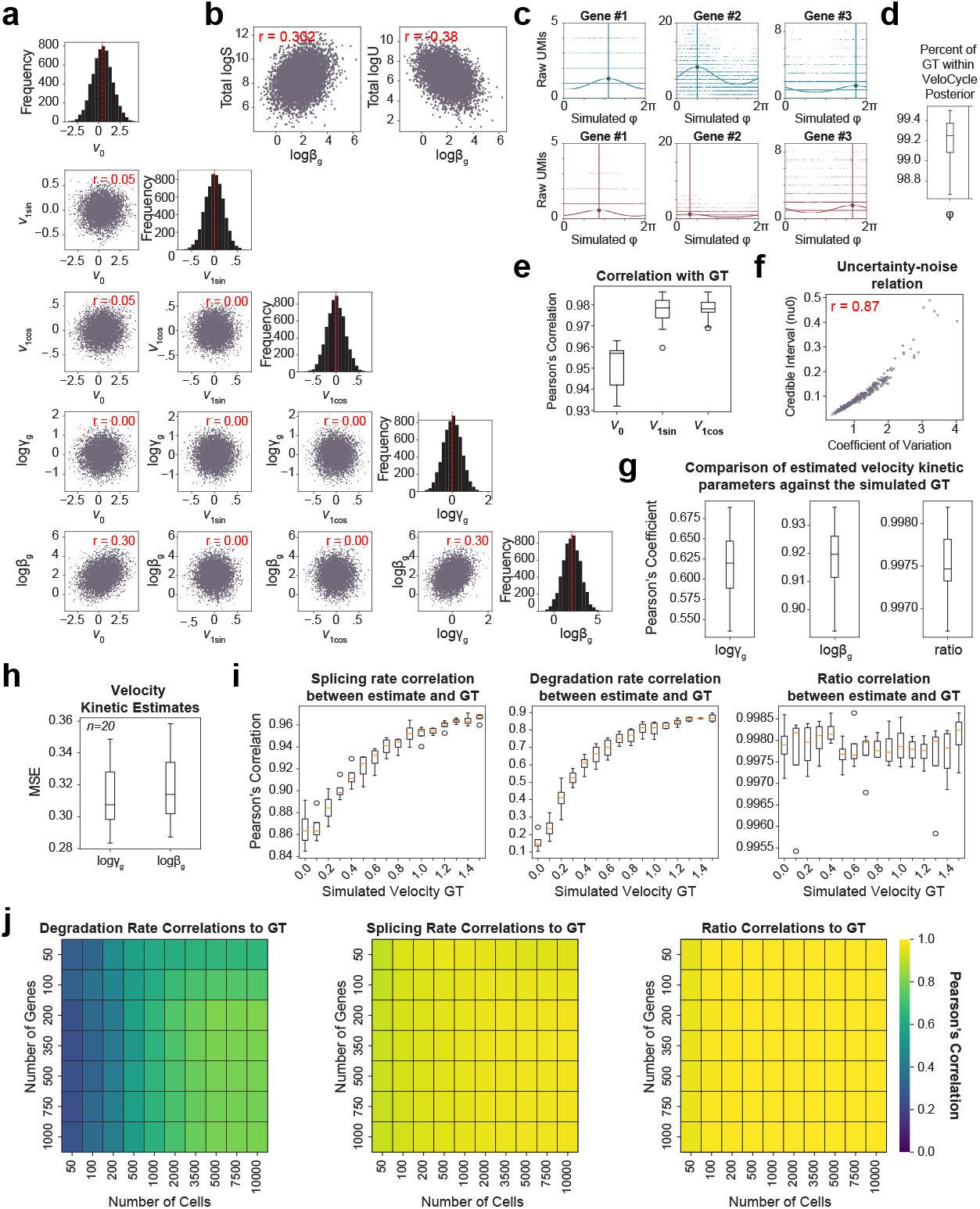
Data generated with structured simulations assists in validation of *VeloCycle*. **(a)** Scatter plot grid of the structured correlation between gene harmonics coefficients (ν_0_, ν_1sin_, ν_1cos_) and kinetic parameters (logβ_g_, logγ_g_) in simulated ground truth (GT) data. Histograms (on diagonal) of the frequency distribution for each simulated latent variable. **(b)** Scatter plots of the simulated data correlation structure between splicing rate (logβ_g_), total spliced (logS), and unspliced (logU) UMI counts. **(c)** Scatter plots of example simulated gene fits for spliced (top; blue) and unspliced (bottom; red) UMIs. Solid curved lines represent gene fits, and vertical lines indicate the phase of peak expression for each gene. **(d)** Box plot of the percent of GT phases within the posterior uncertainty interval estimated, across 20 independently-simulated datasets each containing 3,000 cells and 300 genes. **(e)** Box plots of the mean circular correlation coefficient, across 300 genes, for estimated gene harmonic coefficients obtained compared to the GT. **(f)** Scatter plot of the gene-wise coefficient of variation (a measure of noise) and the credible interval obtained for the gene harmonic coefficient ν_0_. **(g)** Box plots of the mean Pearson’s correlation coefficient for all genes, across 20 independently-simulated datasets, between the estimated and simulated GT for the degradation rate (logγ_g_), splicing rate (logβ_g_), and velocity kinetic ratio (logγ_g_-logβ_g_). **(h)** Box plots of the mean squared error (MSE) for logγ_g_ and logβ_g_ against the simulated GT for 20 datasets in (g). **(i)** Box plots of the mean Pearson’s correlation coefficient between estimated and GT values for logβ_g_, logγ_g_, and velocity kinetic ratio, for all genes across four independently-simulated datasets with 16 different velocity GT between 0.0 and 1.5. **(j)** Heatmaps showing the correlation between estimated and GT values for logβ_g_, logγ_g_, and velocity kinetic ratio using varying numbers of cells and genes. Pearson’s correlation coefficients (red) are indicated in each scatter plot of (a), (b), and (f). Each purple dot represents a single gene in (a), (b), and (f).

**Figure S3.**
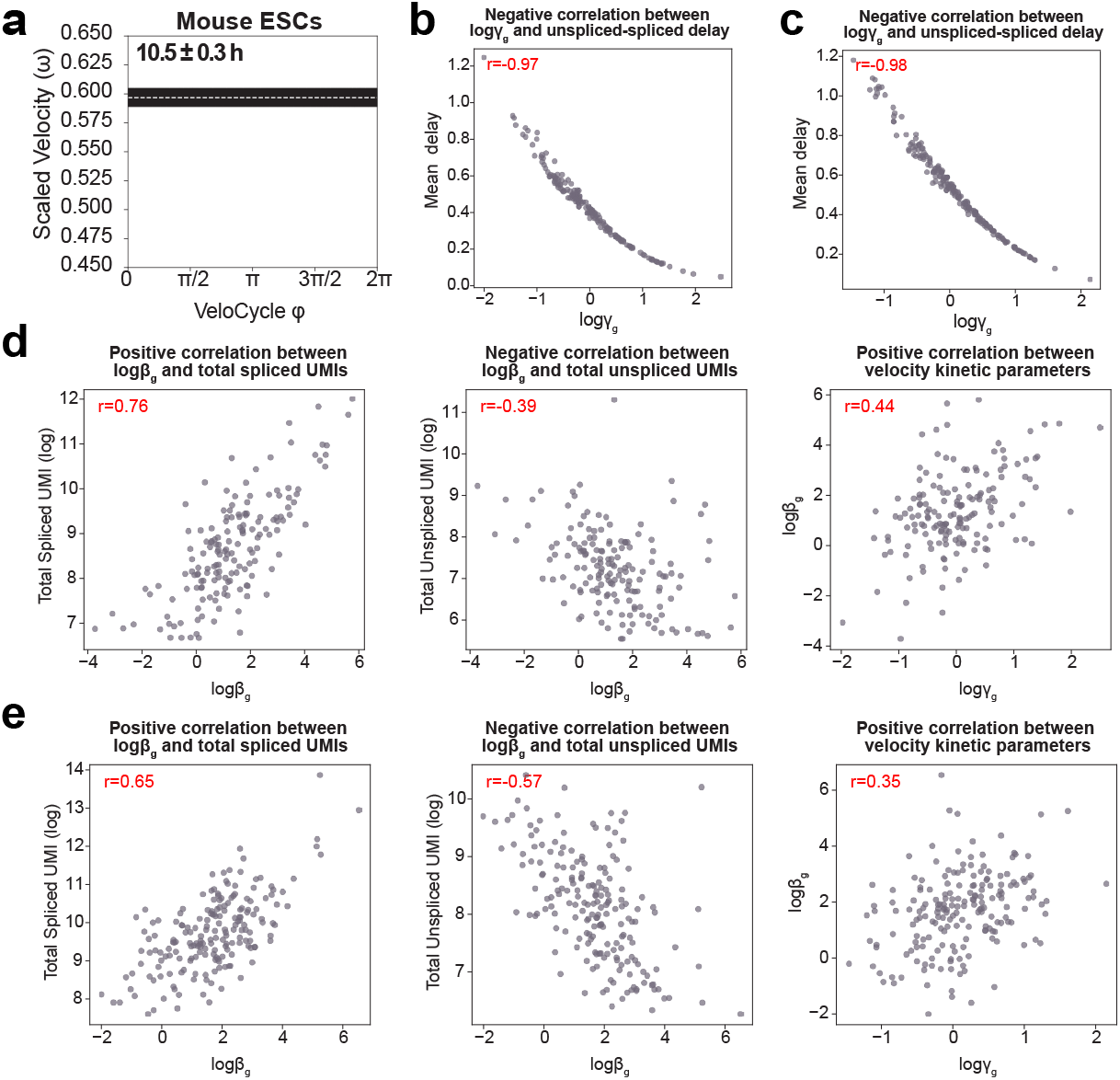
*VeloCycle* accurately measures phase and speed of the cell cycle across species. **(a)** Posterior estimate plot of cell cycle velocity inferred with *velocity-learning* on mouse ESCs [32]. White dashed lines represent the mean of 500 posterior samples; black bars indicate the full posterior interval. **(b)** Scatter plot of the relationship between degradation rate (logγ_g_) and the average unspliced-spliced delay in human fibroblasts. **(c)** Scatter plot of the relationship between degradation rate (logγ_g_) and the average unspliced-spliced delay in mouse ESCs. **(d)** Scatter plots of the relationships among splicing rate (logβ_g_), degradation rate (logγ_g_), and total UMI counts (spliced and unspliced) in human fibroblasts. **(e)** Scatter plots of velocity kinetic parameter relationships for mouse ESCs, as in (d). Pearson’s correlation coefficients (red) are indicated in the top right of plots in (b-e).

**Figure S4.**
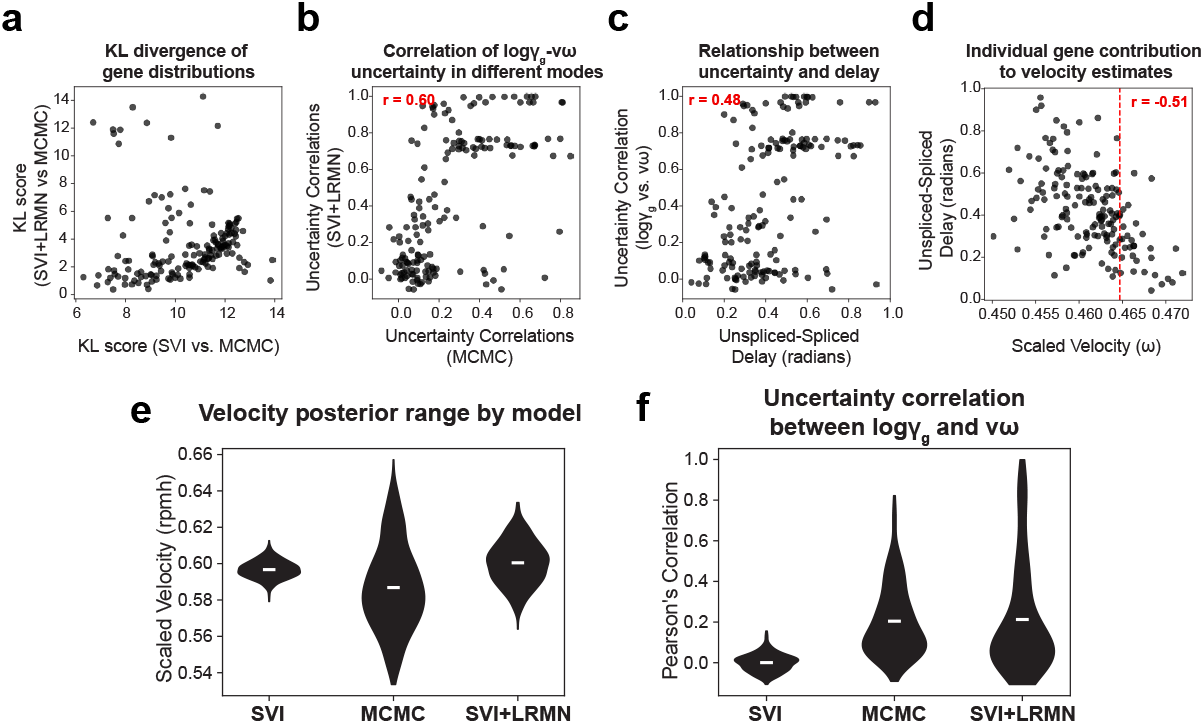
A structured variational distribution yields better velocity uncertainty estimates and reveals relationships among gene kinetic parameters. **(a)** Scatter plot of gene-wise Kullback–Leibler (KL) divergence comparing uncertainty distributions between SVI and MCMC (x-axis) and SVI+LRMN and MCMC (y-axis). A lower KL divergence indicates a greater overlap between the two distributions. **(b)** Scatter plot between the gene-wise logγ_g_-νω uncertainties computed from the posterior of MCMC or SVI+LRMN. **(c)** Scatter plot between unspliced-spliced peak expression delay (radians) and the logγ_g_-νω uncertainty correlation, both obtained using the SVI+LRMN velocity model. **(d)** Scatter plot between the scaled velocity and the unspliced-spliced delay for a leave-one-out estimation approach with *VeloCycle*. Each dot is positioned on the x-axis at the velocity estimate obtained when removing a particular gene (n=160) from the gene set. Each dot is positioned on the y-axis at the position of the unspliced-spiced delay (in radians) for that removed gene. **(e)** Violin plots of the scaled velocity for mouse embryonic stem cells (mESCs), comparable to Fig. 5c. **(f)** Violin plots of the Pearson’s correlations between the degradation rate (logγ_g_) and angular speed (νω) posteriors across all 189 genes for mESCs, comparable to Fig. 5d. Pearson’s correlation coefficients are indicated in red in (b), (c), (d).

**Figure S5.**
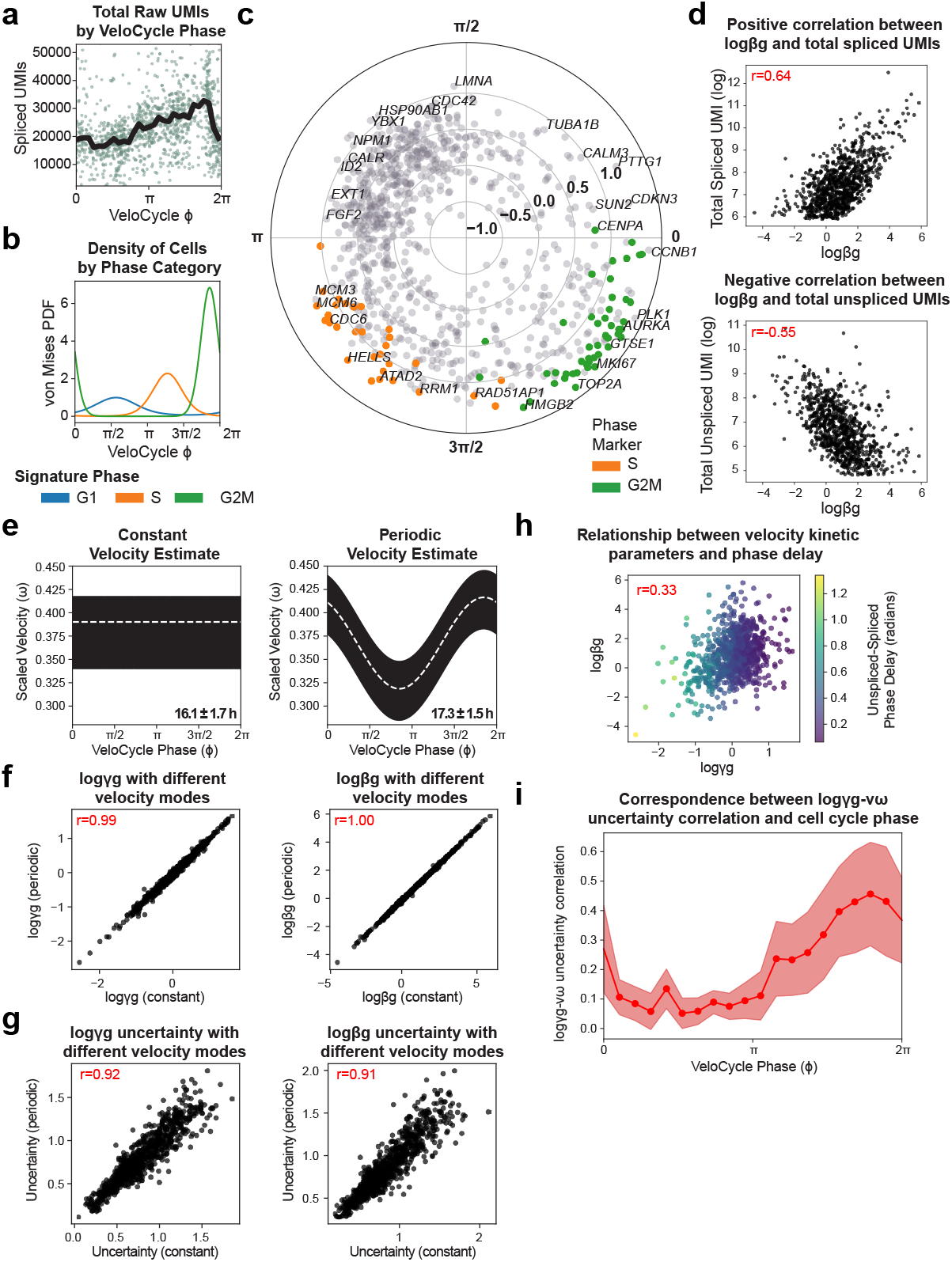
*VeloCycle* coupled with live-cell imaging enables experimental validation of cell cycle speed. **(a)** Scatter plot of total raw spliced UMI counts by continuous cell cycle phase estimated with *VeloCycle* for 1,222 dermal human fibroblasts (dHFs). Black line indicates the binned mean. **(b)** Probability density plot along the *VeloCycle* phase estimation for cells in (a) stratified by categorical phase assignment (G1, S, and G2M) obtained with scanpy. **(c)** Phase space polar plot indicating the phase of peak expression and amplitude for 876 cycling genes used to learn the manifold for dHFs in (a). Each dot represents a gene; genes colored orange (S) or green (G2M) represent marker genes used in traditional categorical cell cycle phase assignment. **(d)** Scatter plots of the relationship between splicing rate and total spliced counts (top) and splicing rate and total unspliced counts (bottom) on a gene-wise basis. **(e)** Posterior estimate plot of constant (left) and periodic (right) velocity estimates obtained for data in (a) using a medium-sized gene set [30]. **(f)** Scatter plots of degradation rates (left) and splicing rates (right) obtained using either the constant (x-axis) or periodic (y-axis) models of velocity estimation. **(g)** Scatter plots of degradation rate (left) and splicing rate (right) posterior uncertainty obtained from 500 posterior samples using either the constant (x-axis) or periodic (y-axis) models. **(h)** Scatter plot of the degradation and splicing rates obtained with the SVI+LRMN model on data in (a-f). Gene-wise dots are colored by the unspliced-spliced phase delay. **(i)** Binned plot of the Pearson’s correlation coefficients between gene-wise degradation rate and velocity posterior uncertainties on dHFs using the SVI+LRMN model of VeloCycle. Pearson’s correlations coefficients are indicated in red text in (d-g).

**Figure S6.**
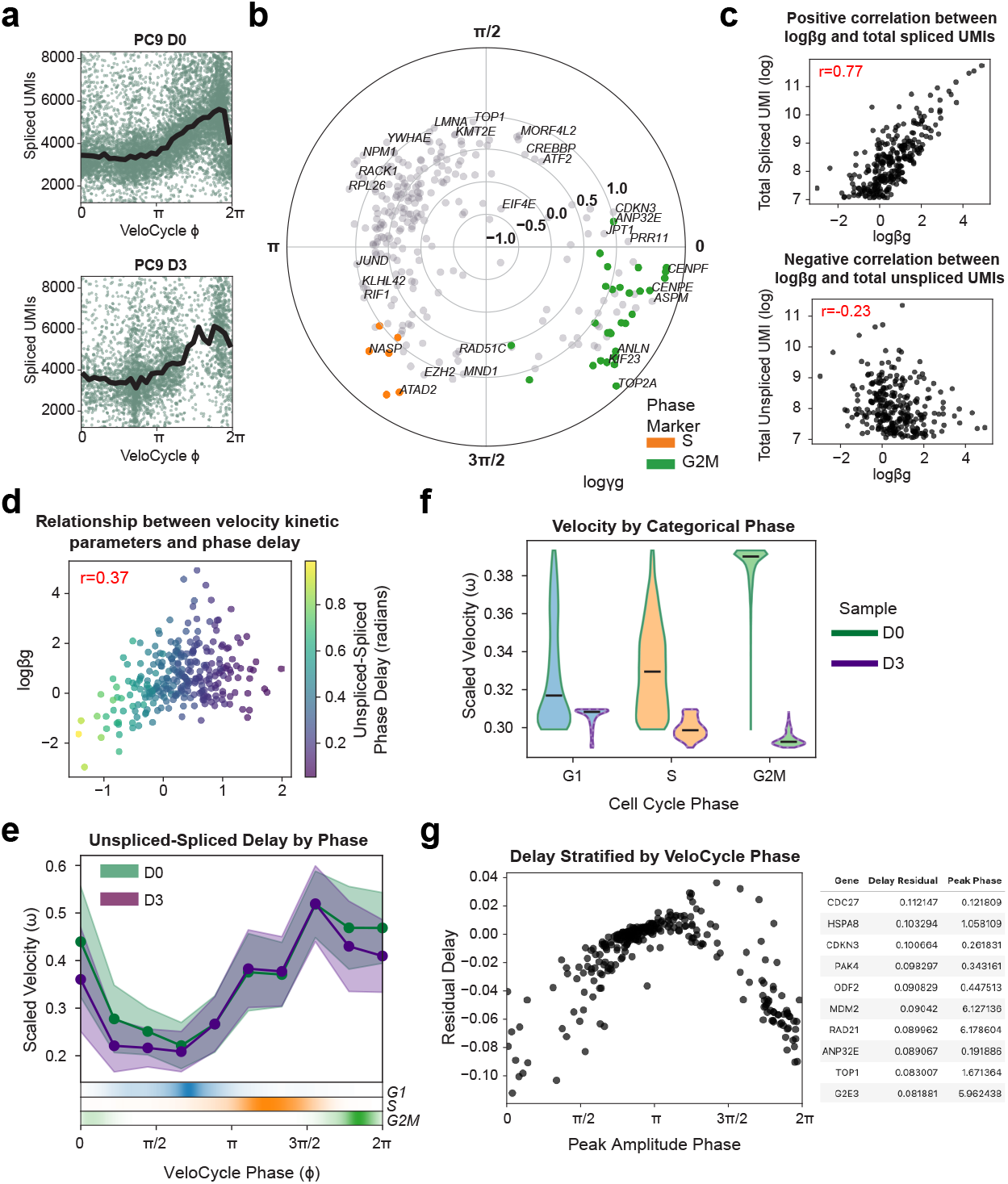
Statistical credibility testing of RNA velocity estimates enables characterization of the effect of erlotinib treatment on lung adenocarcinoma cell line treatment. **(a)** Scatter plot of total raw spliced UMI counts by continuous cell cycle phase estimated with *VeloCycle* for PC9 lung adenocarcinoma cell line populations before (top; D0; 9,927 cells) and after (bottom; D3; 3,943 cells). Black line indicates the binned mean. **(b)** Phase space polar plot indicating the phase of peak expression and amplitude for cycling genes used to learn the manifold for cells in (a). Each dot represents a gene; genes colored orange (S) or green (G2M) represent marker genes used in traditional categorical cell cycle phase assignment. **(c)** Scatter plots of the relationship between splicing rate and total spliced counts (top) and splicing rate and total unspliced counts (bottom) on a gene-wise basis. **(d)** Scatter plot of the degradation and splicing rates obtained with the SVI+LRMN model. Gene-wise dots are colored by the unspliced-spliced phase delay. **(e)** Gene-binned delay between maximum unspliced-spliced expression (in radians) for D0 and D3 samples. **(f)** Violin plots of scaled velocity estimates for D0 and D3, stratified by categorical cell cycle phase. Black horizontal lines indicate the mean. **(g)** Left: scatter plot of peak gene amplitude and residual unspliced-spliced delay (D3-D0) for 273 genes. Right: top 10 differentially delayed genes in D0 versus D3.

**Figure S7.**
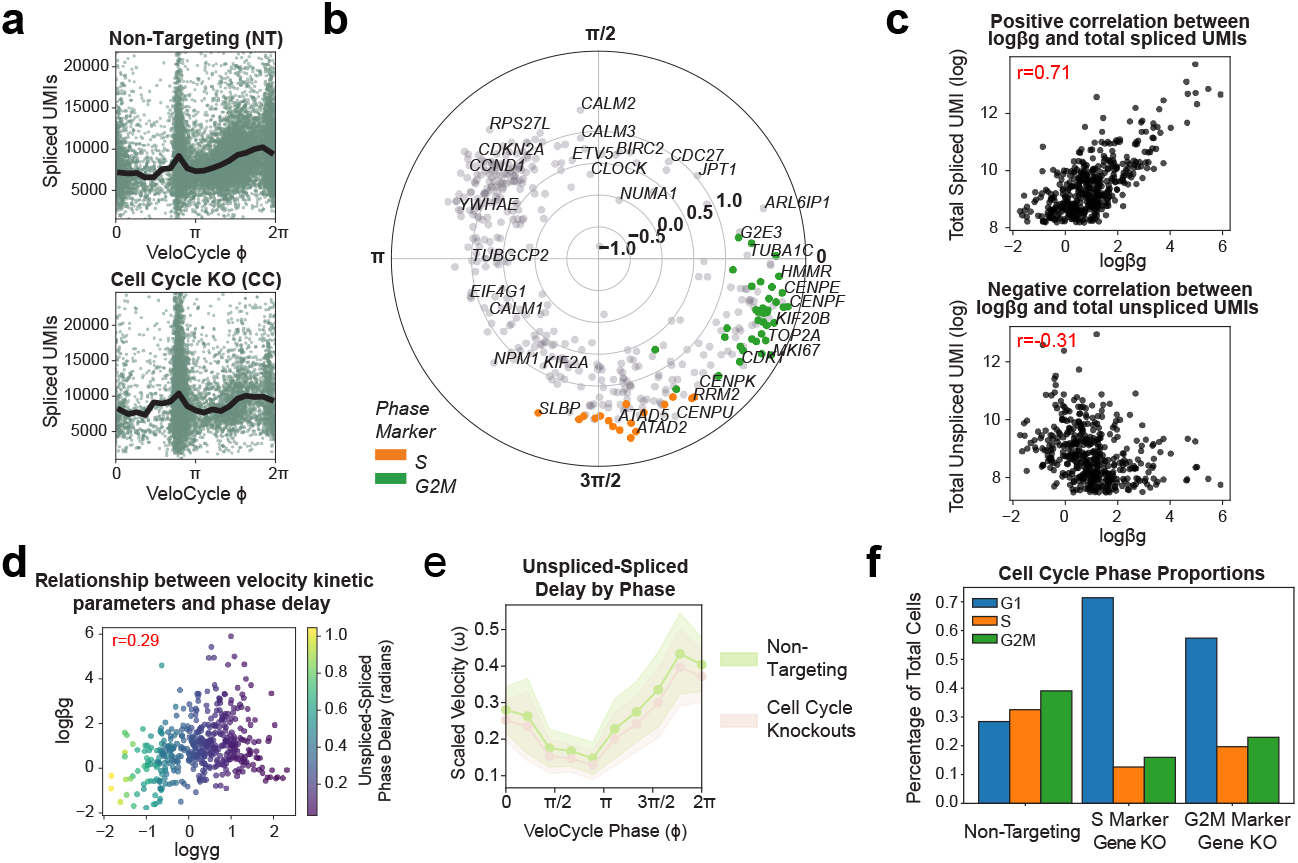
Cell cycle velocity estimation on non-targeting and grouped cell cycle knockout stratifications of RPE1 cells following genome-wide Perturb-seq. **(a)** Scatter plots of total UMIs along the manifold-learning cell cycle phase for non-targeting (NT; top) and cell cycle knockout (CC) strata of genome-wide Perturb-seq data from Fig. 8. **(b)** Phase space polar plot indicating the phase of peak expression and amplitude for 426 cycling genes used to learn the manifold for cells in (a). **(c)** Scatter plots of the relationship between splicing rate and total spliced counts (top) and splicing rate and total unspliced counts (bottom) on a gene-wise basis. **(d)** Scatter plot of the degradation and splicing rates; gene-wise dots are colored by the mean unspliced-spliced phase delay. **(e)** Gene-binned delay between maximum unspliced-spliced expression (in radians) for NT and CC samples. **(f)** Bar plots of categorical cell cycle phase proportions as a percentage of total number of cells, stratified by Perturb-seq non-targeting, S phase marker gene, and G2M marker gene conditions.

## METHODS

### 1. Model specifications for manifold-constrained rna velocity

Gene expression measurements as obtained by scRNA-seq provide a high dimensional snapshot of a cell’s state, with typically *n* ≃ 10^4^ genes being expressed in a cell, of which several thousands are experimentally detected per cell by a nonzero read count. Here, we use the notation *Y*_*c*_ = (*U*_*c*_, *S*_*c*_) for the measurements, with *U*_*c*_ for the unspliced and *S*_*c*_ for the spliced RNA levels (counts), with *S*_*c*_, *U*_*c*_ ∈ ℕ^*n*^.

#### 1.1. The manifold

Many biological processes of interest, such as the cell cycle or a differentiation event, unfold on low-dimensional manifolds *ℳ*. Here, we will consider a parametric representation for *ℳ*: *x* ↦ *s*(*x*) ∈ *ℳ* where *x* is latent coordinate for each cell *c*. Moreover, we will choose the manifold topology based on the biological structure of the problem. For example, given a periodic process such as the cell cycle, we will take *x* ∈ *S*_1_. Typically, the manifold dimension *m* ≪ *n* will be small, and in the case of the cell cycle *m* = 1. As we discuss later, we will learn the function *s*(*x*) from the data (which we will refer to as the *manifold-learning* procedure).

#### 1.2. Measurements and noise model

Measurements for each cell *c* will be linked to the corresponding locations on *ℳ* via realistic noise models. In the case of scRNA-seq, relevant noise models consist of negative binomial (NB) distributions, so that *Y*_*gc*_ *∼* NB(y_g_(x_c_), *α*_g_), with *y*_*g*_(*x*_*c*_) = *E*[*Y*_*gc*_] = (*s*_*g*_(*x*_*c*_), *u*_*g*_(*x*_*c*_)) and 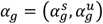. Note that we are assuming for simplicity that *α*_*g*_ is independent of *x* (but this can be relaxed at the expense of an increased number of parameters). This allows us to formulate a likelihood model for the data and approach inference using Bayesian or variational inference.

#### 1.3. RNA velocity and chemical kinetics

In the high-dimensional gene expression space, we expect a rate equation describing the RNA velocity 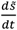 depending both the expectation of spliced and unspliced RNA counts:

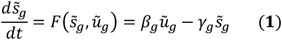

with time-dependent locations 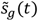 and ũ_*g*_(*t*) and gene-dependent RNA splicing and degradation rates *β*_*g*_ and *γ*_*g*_. Note that here we do not include a corresponding equation for 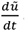 as it will not be needed for the application to the cell cycle. Also, *F* is not explicitly time-dependent and the rates are taken as constants (which could however be relaxed, see below).

#### 1.4. Latent-space dynamics

The key assumption in our approach is that there exists an autonomous (and here deterministic) equation for the dynamics of *x*(*t*) :

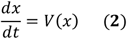

which provides a low-dimensional approximation of the full dynamics (Eq. 1), and that 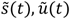 are timedependent through *x*(*t*) :

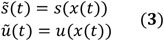

*V*(*x*) is the vector field describing the dynamics in the low dimensional latent space.

#### 1.5. Manifold-constrained RNA velocity

We can now link Eqs. (1), (2) and (3) to obtain

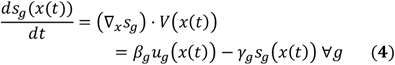

where we have introduced the gene index *g* for clarity and applied the chain rule. *β*_*g*_ and *γ*_*g*_ are the gene-specific splicing and degradation rates.

Eq. (4) provides the basis of our approach as it connects the topology of the low dimensional manifold on the left-hand side with the biology on the right-hand side. Of note, the parameters governing gene dynamics (*β, γ*) could in principle depend on *x* as well.

#### 1.6. Geometric interpretation

By construction, we see that the RNA velocity vector 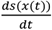 lies in the tangent space of *ℳ* at every point of a trajectory *s*_*g*_(*x*(*t*)). Indeed ∇_*x*_*s* forms an *m*-dimensional basis of the tangent space at each point and *V*(*x*(*t*)) forms the components of the velocity vector in that basis.

#### 1.7. *u*(*x*) and inference

Eq. (4) can also be viewed as specifying *u*(*x*) given *s*(*x*), *V*(*x*), and the parameters *β* and *γ*. This will become central in the implementation. In essence, the optimization algorithm to identify *V*(*x*) and *γ* and *β* coefficients (or functions if we would allow *γ* = *γ*(*x*), etc) such that the predicted RNA velocity 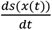 (which lies in tangent space over the entire manifold *ℳ*) is closest to that implied by chemical kinetics and the data *Y*_*c*_ = (*U*_*c*_, *S*_*c*_).

#### 1.8. Duration of biological processes

A benefit of this formulation is that it becomes accessible to estimate the actual duration of biological processes from the trajectories and *V*(*x*) :

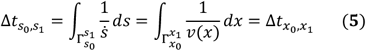

where 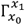 is the trajectory *x*(*t*) that connects the two points *x*_0_ and *x*_1_, and where we have used the change of trajectory variable *s*(*x*). For example, we will be able to estimate cell cycle periods. Moreover, this estimate is by construction independent of the parametrization of the low dimensional manifold.

### 2. Manifolds with *S*1 topology: the cell cycle

Here, we assume that *ℳ* is topologically a circle and therefore we write the coordinate *x* as *φ* ∈ *S*^1^. The equation of the dynamics (Eq. 4) becomes

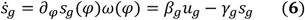

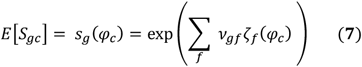

where we assume that *β*_*g*_ and the *γ*_*g*_ are constant along the cell cycle. Of note is that the values of those parameters are constrained by the biology (see section 4 below), which we will enforce through appropriate priors.

*S*^1^ is convenient since it allows use of Fourier series to parameterize the various functions: *s*(*φ*), *u*(*φ*), *ω*(*φ*). Typical cell cycle genes exhibit profiles that can be described by only few harmonics; thus, we will consider up to *k* Fourier components in our expansion (in practice we will by default use one harmonic). Moreover, since *s*(*φ*) is positive, we will use the notation

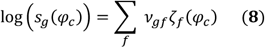

with

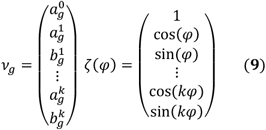

Here *ν*_*g*_ is the vector of gene Fourier parameters written with real numbers.

Using the chain rule, we obtain *u*(*φ*):

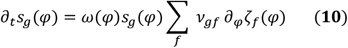

which leads to

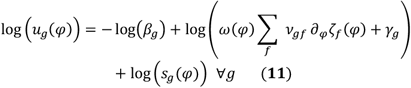

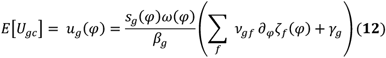

For *ω*(*φ*) we will also be using a Fourier series, limiting ourselves to either constant *ω* or *ω*(*φ*) functions with one harmonic.

#### 2.1. Likelihoods

As explained above with the expressions for *u*(*φ*) and *s*(*φ*), we can calculate a likelihood for the count data over all cells {*Y*_*c*_} = {(*U*_*c*_, *S*_*c*_)}(2.2). To simplify the implementation, we approximate the full joint likelihood for {(*U*_*c*_, *S*_*c*_)} as a product of two factors:

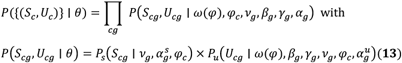

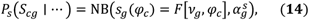

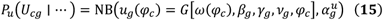

where *θ* is a generic notation for parameters, and *F*[…], *G*[…] show the dependencies of *s*_*g*_, *u*_*g*_ on the other quantities.

We combine these likelihoods with a set of priors into a full Bayesian model (see below) to estimate the joint posterior of *θ*. As indicated above, in our current implementation we simplify the problem by taking two steps: first, we optimize *P*_*s*_ to estimate the cell phases {*φ*_*c*_} and Fourier coefficients {*ν*_*g*_}. We call this step the *manifold-learning* procedure. The second step optimizes *P*_*u*_ and is called *velocity-learning*, using the posterior expectations for 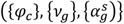 obtained during *manifold-learning* to estimate the remaining quantities 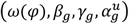.

### 3. Bayesian Model formulation for VeloCycle

Our model includes a mix of biologically defined priors with Empirical Bayes-style priors determined from the data. Our goal will be to estimate an approximation of the joint posterior probability distribution, based on the above expression of the likelihoods:

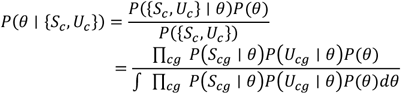

We specify the following priors *P*(*θ*).

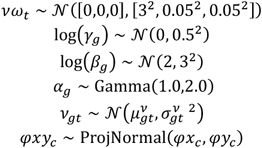

Setting by empirical Bayes the following parameters:

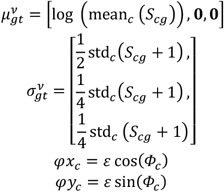

where Φ_*c*_ is obtained from the two first principal components (*w*_1*c*_, *w*_2*c*_) renormalized between [−0.5, 0.5] and computing Φ_*c*_ = tan^−1^ (*w*_2*c*_, *w*_1*c*_). Rotational invariance (e.g., arbitrariness of the first cell *c*0 so that Φ_*c*0_ = 0) is obtained by finding the global phase shift maximizing corr (Φ_*c*_, ∑_*g*_ *S*_*cg*_). The concentration parameter of the projected normal *ε* is set to 5 by default, but can be adjusted depending on the overall confidence in the data quality.

#### 3.1. Variational Distribution - SVI

The variational distribution we use in the base model is mean-field, with marginals of either Normal or Dirac Delta distributed. Specifically

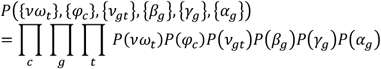

The variational distribution is parametrized as follows (^∧^ indicates the parameters):

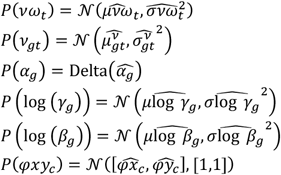

#### 3.2 Variational Distribution - LRMN

The Low Rank Multivariate Normal (LRMN) model considers a variational distribution parametrized to mimic the correlative structure observed between the joint posteriors sampled by Markov Chain Monte Carlo (MCMC) estimation. Specifically, we allow for a covariance and establish specific conditional relationships between the velocity, or angular speed *νω*_*t*_, and the kinetic parameters *β*_*g*_ and *γ*_*g*_. The two main features are: (a) the joint posterior between *γ*_*g*_ and *νω*_*t*_ is parametrized as a low-rank Multivariate Normal, and (b) the marginal posterior of *β*_*g*_ is expressed as conditioned on *γ*_*g*_; namely for each gene *g*, the marginal posterior of *β*_*g*_, through an explicit parameter 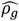, is allowed to correlate with the correspondent *γ*_*g*_.The posterior factorizes as follows:

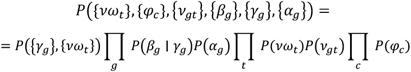

The specific formulation we used is:

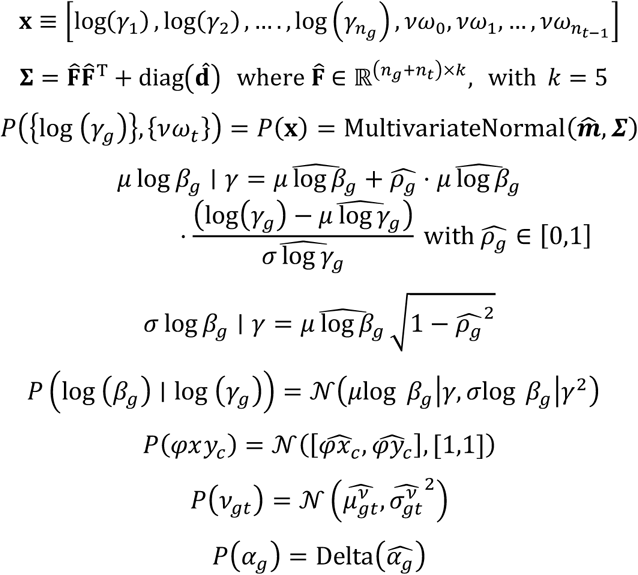

### 4. Model implementation

To estimate an approximation of the joint posterior probability distribution for the angular cell cycle speed (*νω*_*t*_) and the parameters of the *S*^1^ manifold upon which *νω*_*t*_ unwinds, we formulate a likelihood model for the data that we then solve using variational inference in Pyro. This implementation performs estimation of the model latent variables in two steps: *manifold-learning* and *velocity-learning*.

For *manifold-learning*, we estimate the position of each cell along the circular cell cycle manifold (φ) as well as the Fourier series coefficients for each gene (*ν*) describing their periodicity. These variables are then used to model the expectation of log spliced counts (ElogS), which are themselves modeled from the real data and a Negative Binomial. We initialize all variables to the mean of the prior, which is determined using either the first two principal components (φ) or the per-gene mean and standard deviations of spliced expression (*ν*). To allow for differences in average expression levels between different datasets or batches, we also define an offset term (Δ*ν*) for the first gene harmonic coefficient.

For *velocity-learning*, we infer the Fourier coefficients of the angular speed (*νω*) as well as velocity kinetic parameters (*γ* and *β*), conditioned on the mean of the posterior estimates for parameters obtained during *manifold-learning*. These variables are used to model the expectation of log unspliced counts (ElogU), which are themselves modeled from the real data and a Negative Binomial. We initialize all variables to the mean of the prior, which is zero for the angular speed (an assumption of zero cell cycle velocity). In order to enforce positive (*ω*(φ)∑_*f*_ *ν*_*gf*_ ∂_φ_*ζ*_*f*_(*φ*) + *γ*_*g*_) in Eq. (10) during learning, we use a relu function.

Given data, we solve the *VeloCycle* model using stochastic variational inference (SVI) and apply a ClippedAdam optimizer and ELBO loss function, with an evolving learning rate decaying from 0.03 to 0.005 from the first to last training iteration. Typically, we perform 5,000 training iterations for *manifold-learning* and 10,000 training iterations for *velocity-learning*. However, an option to terminating training early is made available, such that no further iterations are executed if the mean loss during the previous 100 iterations is less than 5 units different from the mean loss during the previous 10 iterations.

When performing Monte Carlo Markov Chain (MCMC), we use a No-U-Turn (NUTS) kernel beginning the mean posterior estimates obtained first with SVI. We typically use one chain, 2,000 warm-up sampling steps, and 500 real sampling steps.

*VeloCycle* can be run using either a local CPU or GPU in a few minutes, with significantly improved runtime speeds on GPU, particularly when using a large number of cells (>30,000 cells) or genes (>300 genes). Since there are many more parameters along the gene dimension, scaling up the number of genes reduces runtime more quickly than scaling up the number of cells.

### 4.1. Biological constraints on parameters

Velocity kinetic parameters *β* and *γ* are constrained by the biology. In particular,

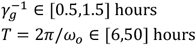

Moreover, the priors for the gene harmonic coefficients are determined for each gene based on the mean level of expression and the variance across all the cells in the data. For the velocity harmonic coefficients, we assume as a prior mean no velocity (i.e., 0) with a wide standard deviation (3.0).

All priors can be easily modified using the velocycle.preprocessing suite of functions and provided to a Pyro model object using the metaparameters (mp) term.

### 4.2. Approximate point estimate for constant cell cycle velocity

To gain an initial insight into the relationship between cell cycle velocity and the expression profiles of the unspliced (*u*)/spliced (*s*) read counts, we used a simplified calculation based on solving the first order differential equation 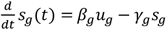, where the degradation rate *γ*_*g*_ is a gene-dependent constant. If we assume that *u*_*g*_(*t*) follows a periodic function with a single harmonic, i.e. *u*_*g*_(*t*) = *u*_0*g*_ (1 + *ε*cos (*ωt* − *φ*_0*g*_)), then *s*_*g*_(*t*) has the same functional form but with a scaled amplitude and shifted phase, depending on the half-life: *s*_*g*_(*t*) = *s*_0*g*_ (1 + *ε*^′^cos (*ωt* − *φ*_1*g*_)), with *ε*^′^ = *ε*cos (Δ*φ*_*g*_), Δ*φ*_*g*_ = (*φ*_1*g*_ − *φ*_0*g*_) and tan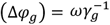. Here, *ω* represents the cell cycle velocity.

Assuming now that we have multiple conditions (or replicates) *c* and that the half lives 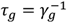 are condition-independent, we observe that the relation

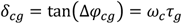

is a rank-1 decomposition of the matrix *δ*_*cg*_, which can be computed using the singular value decomposition (SVD), i.e., *δ*_*cg*_ = *u*_*c*_*dv*_*g*_ + higher rank terms, using standard notation. This allow us to express the condition-specific cell-cycle velocity *ω*_*c*_ in units of inverse mean half lives (noted 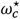)as

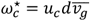

where 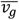 stands for the mean over genes. The cycle-cycle period in units of mean half lives is then 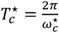.

#### 4.3. Gene sets and quality control filtering

To select genes for velocity analysis that are expected to behave periodically with the cell cycle, we applied one of three differently-sized, literature-based cycling gene sets: “small” containing 97 genes [25], “medium” containing 218 genes [30], and “large” containing 1,918 genes [32]. *VeloCycle* uses the “medium” gene set as a default, in order to minimize the influence of noisy or lowly-expressed genes on manifold and velocity estimation; however, we also employed the “large” gene set in contexts where the sequencing depth and dataset quality are particularly high. The command velocycle.utils.get_cycling_gene_set() can be used to access these human and mouse gene sets. Additional gene filtering based on mean detection of spliced and unspliced counts was also performed as described in the sections below.

#### 4.4. Categorical and continuous cell cycle phase assignment

Categorical cell cycle phase assignment (G1, S, G2/M) was performed using the scanpy function sc.tl.score_genes_cell_cycle(), as previously described [25]. Continuous cell cycle phase assignment using DeepCycle on both simulated and real datasets was achieved using the velocity information obtained from scvelo.pp.moments [4] and standard parameters described in the original publication [30].

#### 4.5. Inference of the unspliced-spliced delay

To compute the unspliced-spliced delay from the results of *VeloCycle*, we calculated the difference between phases of peak expression of unspliced and spliced UMIs on a per gene basis (in radians) using the estimated expectations of unspliced (ElogU) and spliced (ElogS) counts.

#### 4.6. Posterior probability sampling

Unless otherwise stated, the latent variables and associated estimate uncertainties were collected from 500 posterior samples after model training using pyro.infer.predictive, and credibility intervals were measured between the 5th and 95th percentiles.

Estimates for the cell cycle velocity obtained from the velocity function ω(φ) were scaled by the mean degradation half-life, i.e., mean(γ_g_). To infer the cell cycle period over the entire cell cycle, we sampled from the velocity function on a grid of 20 phases (from 0 to 2π) and took the area under the curve using scipy.integrate.trapz. The posterior mean, 5th percentile, and 95th percentile were then computed using numpy.mean and numpy.percentile. The full uncertainty range of the posterior estimate was computed by taking the difference between the 95th and 5th percentile estimates.

### 5. Structured data simulations and sensitivity analyses of *V**elo**C**ycle*

To properly validate the performance of *VeloCycle* on datasets with a ground truth for all latent variables of the *manifold-learning* and *velocity-learning* procedures, we employed a new structured simulation approach to preserve relationships among velocity kinetic parameters (splicing rate β, degradation rate γ) and gene harmonics (*ν*_0_, *ν*_1sin_, *ν*_1cos_). These relationships are expected in real data [2] and are necessary in simulations to avoid improbable scenarios where the ratio of unspliced to spliced counts is unrealistically high or low. We expect that genes containing more velocity information should be those with a larger unspliced-spliced delay and slower splicing and degradation rates; genes with too fast kinetics will provide limited signal in scRNA-seq data. Thus, we formulated a generative *VeloCycle* model that imposes a correlation structure among the gene harmonic and velocity kinetic rate parameters for the sole purpose of sampling simulated data (and not for use during inference itself). We defined correlations as follows: a weak positive correlation among the gene harmonic coefficients (r=0.05; assuming only one sine and cosine term per gene), a moderate positive correlation between the splicing rate and zeroth gene harmonic coefficient *ν*_0_ (r=0.30), and a moderate positive correlation between splicing and degradation rates (r=0.30).

Using this correlation matrix, simulated datasets were generated by randomly sampling for a user-defined number of genes and cells, from a pyro.dist.MultivariateNormal. These variables, along with a user-defined ground-truth cell cycle speed (νω) and a cell-specific phase (φ) sampled from a random uniform distribution between 0 and 2π, were plugged into the velocity equations to compute an expectation for unspliced (ElogU) and spliced (ElogS). Finally, raw data (S and U) was sampled from a pyro.dist.GammaPoisson using the expectations and a noise parameter (shape_inv) sampled from pyro.dist.Gamma. All simulated data generated for this study are available on Zenodo (see **Data Availability**). Additional datasets can be simulated using the velocycle.utils.simulate_data function.

Evaluation of the *manifold-learning* step was performed using 20 datasets, each containing 3,000 cells and 300 genes, independently simulated with a ground truth velocity of 0.4. The same datasets were also used for validation of the *velocity-learning* step. To perform sensitivity analysis on the number of cells and genes, four independently simulated datasets containing 10,000 cells and 1,000 genes were generated; data subsets were used to test the model’s performance on varying numbers of cells (from 100 to 5,000 for *manifold-learning* and from 50 to 10,000 for *velocity-learning*) and genes (from 100 to 1,000 for *manifold-learning* and from 50 to 1,000 for *velocity-learning*). To assess *velocity-learning* performance on datasets with different ground truth velocities, we simulated four datasets with shared kinetics and gene harmonic parameters, but one of 16 different ground truth velocities from 0.0 to 1.5.

Circular correlations between estimated and simulated ground truth variables were computed using velocycle.utils.circular_corrcoef, which converts the input data into unit circle coordinates and computes a correlation by finding the mean of the product of estimated values and the complex conjugate of the ground truth values. To compare phases obtained with *VeloCycle* to those from DeepCycle, the same simulated datasets were used to compute velocity moments with sc.pp.moments followed by running DeepCycle with default parameters described in the original publication [30].

### 6. *V**elo**C**ycle* estimation across multiple standard scRNA-seq datasets

In this work, we performed manifold geometry and cell cycle velocity estimation with *VeloCycle* on a number of published datasets from different technologies, species, and sampling contexts. For all datasets, the original raw data were re-processed using *velocycle* [2] to obtain spliced and unspliced count matrices. A general procedure for running *VeloCycle* on these types of scRNA-seq data has been described above and is supported by tutorials on the corresponding GitHub page for these works. Here, we provide a summary of any specific filtering criteria and parameters used on a dataset-dependent basis.

#### 6.1. FACS-sorted mouse embryonic stem cells [31]

*VeloCycle* estimation of cell cycle phases and gene harmonics was performed on 279 single cells from a culture of Smart-seq2 mouse embryonic stem cells (mESC) using the standard parameters. Genes used in *manifold-learning* were those from the “large” gene set (Gene Ontology; 1,918 genes) available in velocycle.utils, after filtering out genes with <=0.5 mean unspliced counts per cell or with <=2 mean spliced counts per cell (1,358 genes remaining). *Manifold-learning* was performed using 3,000 training steps.

To evaluate the predictive capacity of categorical cell cycle phase (G1, S, or G2/M) using the *VeloCycle* phases, a DecisionTreeClassifier from sklearn.tree was trained with 65% of cells, reserving 35% of cells as a test set and for calculation of a confusion matrix. To compare with a model using the total gene expression matrix to predict categorical cell cycle phases, the linear LogisticRegressionCV model from sklearn.linear_model was trained (cv=5) using the same train-test cell split as with the decision tree.

#### 6.2. Mouse embryonic stem cells and human fibroblasts [30]

*VeloCycle* was run separately on 5,191 single cells from a culture of mouse embryonic stem cells (mESC) and on 2,557 single cells from a culture of human fibroblasts using the standard parameters. Non-cycling cells were filtered out prior to analysis according to the author’s annotations. Genes used in *manifold-learning* were those from the “medium” gene set (DeepCycle; 218 genes) available in velocycle.utils, after filtering out genes with <=0.1 mean unspliced UMIs per cell or with <=0.3 mean spliced UMIs per cell (189 genes and 160 genes remaining for mESC and fibroblasts, respectively). *Manifold-learning* was performed using 5,000 training steps, and *velocity-learning* was performed using the “normal” guide and the constant-velocity model for 10,000 training steps. Comparisons to DeepCycle phases were made using the published estimates described for these exact datasets in the original study [30].

#### 6.3. Human dermal fibroblasts [39]

*VeloCycle* was run on 1,222 single cells from a culture of untreated dermal human fibroblasts (dHFs) using the standard parameters; non-cycling cells were excluded using the author’s annotations. Genes used in *manifold-learning* were those from the “large” gene set (Gene Ontology; 1,918 genes) available in velocycle.utils, after filtering out genes with <=0.1 mean unspliced UMIs per cell or with <=0.3 mean spliced UMIs per cell (876 genes remaining). *Manifold-learning* was performed using 5,000 training steps, and *velocity-learning* was performed with both the constant-velocity and periodic-velocity models for 10,000 training steps using the low-rank multivariate normal (“lrmn”) guide.

Time-lapse microscopy data, including cell segmentation and tracking, for dHFs was obtained from the originally-published study and is available on the corresponding Zenodo page: 10.5281/zenodo.6245943. A cell was determined to be poorly-tracked and excluded from analysis if it had a measured cell cycle length less than 8 hours or greater than 32 hours.

#### 6.4. PC9 lung adenocarcinoma cell line [42]

*VeloCycle* was run jointly on data from PC9 lung adenocarcinoma cell line prior to (D0: 7,927 cells) and after (D3: 3,743 cells) treatment with erlotinib using the standard parameters. Genes used in *manifold-learning* were those from the “large” gene set (Gene Ontology; 1,918 genes) available in velocycle.utils, after filtering out genes with <=0.1 mean unspliced UMIs per cell or with <=0.1 mean spliced UMIs per cell. After an initial *manifold-learning* step, only genes with a Pearson’s correlation between the unspliced and spliced counts >=0.8 and a predicted unspliced-spliced delay greater than >=-0.25 were retained. *Manifold-learning* was performed using 5,000 training steps, and *velocity-learning* was performed using the “lrmn” guide and both the constant-velocity and periodic-velocity models for 10,000 training steps.

#### 6.5. Radial glial progenitors from the developing mouse brain [48]

*VeloCycle* was run jointly on all radial glia progenitor cells from the E10 time point, stratified by regional identity (forebrain: 3,293 cells; midbrain: 2,388 cells; hindbrain: 2,012 cells) using the standard parameters. Genes used in *manifold-learning* were those from the “large” gene set (Gene Ontology; 1,918 genes) available in velocycle.utils, after filtering out genes with <=0.05 mean unspliced UMIs per cell or with <=0.1 mean spliced UMIs per cell. After an initial *manifold-learning* step, only genes with a Pearson’s correlation between the unspliced and spliced counts >=0.8 and a predicted unspliced-spliced delay greater than >=-0.10 were retained. *Manifold-learning* was performed using 5,000 training steps, and *velocity-learning* was performed using the “lrmn” guide and the constant-velocity model for 10,000 training steps.

Similarly, *VeloCycle* was run jointly on all radial glia progenitor cells from the E14 and E15 time points, stratified by regional identity (forebrain: 2,460 cells; midbrain: 307 cells; hindbrain: 176 cells) using the standard parameters. With the same gene filtering steps as with the E10 analysis above, 239 genes were used. *Manifold-learning* was performed using 5,000 training steps, and *velocity-learning* was performed using the “lrmn” guide and the constant-velocity mode for 10,000 training steps.

To spatially visualize *VeloCycle* speed estimates at the E10 time point, we ran the BoneFight algorithm to map scRNA-seq clusters to a corresponding spatial transcriptomics dataset of hybridization-based in situ sequencing (HybISS) from the same study, then colored the corresponding clusters by their velocity estimate.

#### 6.6. Genome-wide Perturb-seq RPE1 cells data [58]

To ensure analysis was performed only on RPE1 cells with a complete knockdown of the individual gene target, we filtered out cells containing non-zero unspliced or spliced UMI reads for the targeted gene. *VeloCycle* was run initially on a subset of data in two conditions: (1) the set of control, non-targeting cells (NT: 11,485 cells) and a grouped set of cells where a gene from the “small” cell cycle marker list were targeted for knockdown (CC-KO: 6,275 cells). Genes used in *manifold-learning* were those from the “medium” gene set (DeepCycle; 218 genes) available in velocycle.utils, after filtering out genes with <=0.1 mean unspliced UMIs per cell or with <=0.2 mean spliced UMIs per cell. After an initial *manifold-learning* step, only genes with a Pearson’s correlation between the unspliced and spliced counts >=0.7 and a predicted unspliced-spliced delay greater than >=-0.5 were retained (120 genes remaining). *Manifold-learning* was performed using 5,000 training steps, and *velocity-learning* was performed using the “lrmn” guide and the constant-velocity mode for 10,000 training steps.

To perform stratified analysis on a larger batch of gene knockout conditions, we selected all cells from any conditions represented by more than 75 cells, leaving a total of 167,119 cells and 986 conditions. We then conditioned on the gene harmonic coefficients obtained with the coarser analysis using NT and CC-KO cells, and performed *manifold-learning* for 5,000 training steps to estimate cell cycle phases for all cells and conditions. We then performed *velocity-learning* for 10,000 training steps on the entire dataset, estimating an individual constant velocity for each gene knockout condition.

#### 6.7. RPE1 cells (newly-generated for this study)

To estimate the unspliced-spliced delay and cell cycle velocity between two identical replicates of FUCCI-RPE1 cells (replicate 1: 4,265 cells; replicate 2: 9,994 cells), *manifold-learning* was run on the “medium” gene set available in velocycle.utils, after filtering out genes with <=0.1 mean unspliced UMIs per cell or with <=0.3 mean spliced UMIs per cell (136 genes remaining). *Manifold-learning* was performed using 3,000 training steps, and *velocity-learning* was performed with both the constant-velocity and periodic-velocity modes for 10,000 training steps using the low-rank multivariate normal (“lrmn”) guide.

Likewise, for the third sample of wild-type RPE1 cells (3,354 cells), *manifold-learning* was run on the “medium” gene set available in velocycle.utils, after filtering out genes with <=0.1 mean unspliced UMIs per cell or with <=0.3 mean spliced UMIs per cell (128 genes remaining). *Manifold-learning* was performed using 3,000 training steps, and *velocity-learning* was performed with both the constant-velocity and periodic-velocity modes for 10,000 training steps using the low-rank multivariate normal (“lrmn”) guide.

### 7. Experimental procedures

#### 7.1. Cell culture

FUCCI-RPE-1 cells (see **Fig. 4**), a gift from Battich et al[62] were cultured at 37°C and 5% CO_2_ in DMEM/F12 medium (Gibco 11320033) supplemented with 1% NEAA (Gibco 11140-035), 1% Penicillin/Streptomycin (Sigma Aldrich G6784) and 10% FBS (Gibco 10437-028).

Additional RPE-1 cells (see **Fig. 5**) were obtained from ATCC and cultured at 37°C, 20% O_2_, and 5% CO_2_ in DMEM/F12 medium (Gibco 21331-020) supplemented with 1% MEM NEAA (Sartorius 01-340-1B), 0.5% sodium pyruvate 1% penicillin/streptomycin/glutamine, and 10% FBS. Media was replaced daily and cells were passaged twice a week. RPE-1 cells were maintained in culture for at least two passages and confirmed to be free of mycoplasma.

#### 7.2. scRNA-seq library preparations

For the preparation of scRNA-seq libraries, an experimental setup was designed to mimic the conditions used for live-cell imaging. FUCCI-RPE-1 cells (see **Fig. 4**) were seeded (7000 cells/cm^2^) in duplicate 2 days before collection. On collection day, cells were detached with trypsin, washed with PBS, counted and diluted to a cell concentration of 1000 cells/uL. Barcoded cDNA libraries were generated from single cell suspensions using the 10x Genomics Chromium v3.1 dual-index system. The procedure was done in accordance with the manufacturer’s instructions, with a goal of 4,000 cells per library. Samples were individually indexed and evenly pooled together. After quality control, libraries were sequenced on an Illumina HiSeq4000 platform, with a depth of approximately 300 million reads per sample, by the EPFL Gene Expression Core Facility (GECF).

Similarly, RPE-1 cells (see **Fig. 5**) were detached using Trypsin–EDTA solution A 0.25% (Biological Industries; 030501B) for 5 min at 37°C. Trypsin was neutralized with medium including 10% FBS and cells were centrifuged at 250 rcf for 5 min, followed by washing and resuspension in PBS with 0.04% BSA. The cell suspension was filtered with a 40-μm cell strainer to remove cell clumps. A cell viability percentage higher than 90% was determined by trypan blue staining. Cells were diluted to a final concentration of 700 cells per μl. scRNA-seq libraries were generated using the 10x Genomics Chromium v3.1 dual-index system. The procedure was done in accordance with the manufacturer’s instructions, with a goal of 3,000 cells per library. Samples were then indexed and sequenced on an Illumina NovaSeq 6000 platform by the EPFL Gene Expression Core Facility (GECF).

As with all publicly available datasets, raw fastq files were processed with CellRanger using the default human reference transcriptome to obtain count matrices. To obtain unspliced and spliced count matrices, we used velocyto version 0.17.17.

#### 7.3. Live-image microscopy and cell tracking experiments with RPE1 cells

RPE-1 cells were seeded on glass bottom 6 well chamber slides (IBIDI) to reach 30% confluence after one day. Cells were then imaged on a PerkinElmer Operetta microscope under controlled temperature and CO_2_ every 10.25 minutes using brightfield and digital phase contrast (DPC) with a 10x (0.35 NA) air objective, binning of 2 and the speckle scale set to 0 under non-saturated conditions. Cell division tracking was achieved by stacking time-course images and manually tracing cell movement and division with napari [68]. Between 20-25 RPE1 cells were tracked from 15 different fields of view by three different individuals (A.R.L., A.H., A.V.) for a total of 337 cells used to estimate a ground truth.

#### 7.4. Cumulative EdU and p21 staining experiments

Cells were seeded on poly-L-lys-coated 24 well plates to reach 30% confluence after one day. After a day, 10 μM EdU (Invitrogen - #A10044) was added to the media, and cells were fixed at different time points after EdU addition: 30 min, 1h, 2h, 3h, 5h, 8h, 12h, 24h, 32h, 36h, 49h, 56h, 72h. For each time point, cells were fixed in 4% PFA for 10 minutes, washed twice with PBS, and processed for EdU (5-ethynyl-2’-deoxyuridine) detection according to the manufacturer’s instructions (Click-iT EdU Alexa Fluor 647 Imaging kit from Invitrogen #C10340). Additionally, cells were permeabilized with 0.2% triton and stained overnight at 4°C with p21 Waf1/Cip1 (12D1) Rabbit mAb (Cell Signaling Technology #2947) and revealed with a secondary antibody conjugated to Alexa Fluor-488. After staining, cells were imaged on a Leica DMi8 (20x NA 0.8).

To quantify the signal intensities of p21 and EdU, we segmented nuclei in the DAPI channel using stardist [69]. We obtained the average intensity for both signals per nucleus by sub-setting the corresponding channel using segmentation masks. The intensity of p21 was normalized per image (percentile-based, p_min=1, p_max=99.8), as its intensity profile was expected to be approximately constant in time; conversely, the intensity of EdU was not normalized as it was expected to increase with time. Thresholds were selected observing the (bimodal) signal distribution across nuclei in all timepoints.

First, to compute the time it takes, on average, for a cell to traverse through two consecutive S phases (tEdU), we applied the Nowakowski method [52, 70] on data collected at multiple time points for a total number of 678,204 cells. The Nowakowski method assumes a linear growth of EdU+ cells, until reaching a plateau where all cycling cells are positive for EdU. We obtained a linear fit of the growth and determined the x-value at the intersection between growth and plateau (A), the y-intercept of the linear fit (B), and the y-value of the plateau (GF). With these, we could compute tEdU as follows: tEdU = (B*A)/(GF-B) + A.

However, cells may on occasion exit the cycle to a G0 phase, and then re-enter at a later time [71]. To correct for this, we plotted the fraction of EdU+/p21+ cells among the p21+ population to estimate the G0 duration (tG0). We determined the tG0 to be equal to the time point at which fraction of EdU+/p21+ cells plateaued, after which no statistically significant changes were detected (Tukey’s multiple comparison test). The corrected estimate for the cell cycle duration was finally calculated as: tc = tEdU - (%p21 * tG0), where %p21 corresponded to the mean fraction of p21+ cells.

